# Closure of the γ-tubulin ring complex by CDK5RAP2 activates microtubule nucleation

**DOI:** 10.1101/2023.12.14.571518

**Authors:** Yixin Xu, Hugo Muñoz-Hernández, Rościsław Krutyhołowa, Florina Marxer, Ferdane Cetin, Michal Wieczorek

**Affiliations:** Department of Biology, Institute of Molecular Biology and Biophysics, ETH Zürich, 8093 Zürich, Switzerland

## Abstract

Microtubule nucleation in cells is templated by the γ-tubulin ring complex (γ-TuRC), a 2.3 MDa multiprotein assembly concentrated at microtubule organizing centers (MTOCs). Current γ-TuRC structures exhibit an open conformation that deviates from the geometry of α/β-tubulin in the microtubule, potentially explaining their low in vitro microtubule-nucleating activity. Several proteins have been proposed to activate the γ-TuRC, but the mechanisms underlying activation are not known. Here, we isolated the porcine γ-TuRC using CDK5RAP2’s centrosomin motif 1 (CM1) and determined its structure with cryo-electron microscopy. 3D heterogeneity analysis revealed an unexpected conformation of the γ-TuRC, in which five protein modules containing MZT2, GCP2, and CDK5RAP2 decorate the outer face of the holocomplex. These decorations drive a long-range constriction of the γ-tubulin ring, bringing the GCP2/GCP3-rich core of the complex in close agreement with the architecture of a microtubule. A purified CDK5RAP2 fragment stimulated the microtubule nucleating-activity of the porcine γ-TuRC as well as a reconstituted, CM1-free human complex in single molecule assays. Our results show that CDK5RAP2 activates the γ-TuRC by promoting γ-tubulin ring closure, providing a structural mechanism for the regulation of microtubule nucleation by CM1 motif proteins in mammals and revealing conformational transitions in γ-tubulin that prime it for templating microtubule nucleation at MTOCs.

## Introduction

The nucleation and organization of the microtubule cytoskeleton depends on the γ-tubulin ring complex (γ-TuRC), a multiprotein assembly that provides a polymerization template for microtubules [1–4]. Perturbing the function of this essential complex leads to inhibited axonal growth [1], defective mitotic spindle formation [2], and severe developmental disorders [3–5]. Although a substantial fraction of γ-tubulin in cells is in a soluble form [6], γ-TuRC-mediated microtubule nucleation is usually suppressed in the cytosol. Instead, its activity is restricted to specialized organelles such as centrosomes and other microtubule organizing centers (MTOCs) [7]. Microtubule nucleation in cells is also dynamically regulated; the level of γ-TuRC recruited to prophase centrosomes is roughly three-fold higher than for interphase centrosomes [8]. How cells facilitate correct γ-TuRC recruitment and activation is an open question.

The human γ-TuRC has a molecular weight of ∼2.3 MDa and consists of γ-tubulin, <u>γ</u>-tubulin <u>C</u>omplex <u>P</u>rotein <u>2</u> (GCP2), GCP3, GCP4, GCP5, and GCP6, MZT1, MZT2, and several γ-TuRC-associated proteins [9]. These proteins form an asymmetric cone-shaped structure that organizes a semi-helical ring of γ-tubulin [10–12]. In all available vertebrate γ-TuRC structures [10,11,13–16], the γ-tubulin ring is in an open conformation that approaches - but does not match - the architecture of a 13-protofilament microtubule [17]. This may explain why the γ-TuRC is such a poor microtubule-nucleating factor *in vitro* [11,15,18,19], and suggests that γ-tubulin ring closure is required for efficient microtubule nucleation [11,20,21]. In current models, the γ-tubulins associated with GCP4, GCP5 and GCP6 are more displaced and are proposed to be major sites of allosteric regulation [11,15,22], but the potential mechanisms are not clear.

The local activities of α/β-tubulin as well as microtubule-associated proteins play an important role in regulating templated nucleation [23]. However, accessory proteins which recruit the γ-TuRC to MTOCs and may stimulate its microtubule-nucleating activity have also been identified. The most well-studied candidate γ-TuRC-activating proteins are those containing the centrosomin motif 1 (CM1). This ∼60-70 amino acid motif is typically found at the N-terminus of orthologous centrosomal and spindle pole body proteins such as *D. melanogaster* centrosomin [24], Mto1 in *S. pombe* [25], *S. cerevisiae* Spc110p [26], and *H. sapiens* CDK5RAP2 [27]. In budding yeast, the γ-TuRC is built from repeating units of the so-called γ-Tubulin Small Complex, or γ-TuSC, an evolutionarily conserved subcomplex composed of γ-tubulin, Spc97 (GCP2), and Spc98 (GCP3). Several structures of the CM1 motif from *S. cerevisiae* Spc110p biochemically reconstituted with the γ-TuSC have been determined [19,20,28,29]. These structures suggest that CM1-binding aids in γ-TuSC oligomerization by contributing a key α-helix that bridges across the inter-γ-TuSC interface. However, it is not clear whether CM1-binding alone resolves an open γ-tubulin organization also observed in γ-TuSC helical oligomers [28]. Moreover, budding yeast do not contain GCP4, GCP5, or GCP6, and a high-resolution structure of the intact CM1-bound *S. cereviseae* γ-TuRC is not available, making it difficult to infer a universal molecular mechanism of γ-TuRC recruitment and activation by CM1 motif proteins from this system alone.

Human CDK5RAP2 is a 215 kDa CM1 motif-containing protein localized to centrosomes and the Golgi [30,31]. RNAi depletion of CDK5RAP2 reduces centrosomal microtubule nucleation without interfering with γ-TuRC assembly [27,32]. As is characteristic for many centrosomal proteins, CDK5RAP2 is predicted to be rich in extended α-helices with a high propensity for forming coiled-coils [27,33]. The 57 residue-long N-terminal CM1 motif of CDK5RAP2 contains a sequence that stimulates microtubule nucleation from the γ-TuRC *in vitro* (a.a. 57-88), while another segment auto-inhibits this activating effect (a.a. 96-114) [32,34]. The structure of the activating portion of CDK5RAP2’s CM1 motif has previously been determined in the human γ-TuRC [12]. In this structure, CDK5RAP2 binds a single site in the holocomplex, and forms a dimeric coiled-coil that is positioned between the C-terminal <u>γ</u>-tubulin <u>RI</u>ng <u>P</u>rotein <u>2</u> (GRIP2) domain of GCP2, a subcomplex of MZT2 intercalated with GCP2’s <u>N</u>-terminal α-<u>H</u>elical <u>D</u>omain (GCP2-NHD), and the C-terminal GRIP2 domain of GCP6. While the presence of CDK5RAP2 modifies the local organization of the GCP2- and GCP6-bound γ-tubulins into a more microtubule lattice-like arrangement [28], most of the γ-tubulin ring still displays an open configuration indistinguishable from CDK5RAP2-free γ-TuRCs [10]. It has been proposed that CDK5RAP2 could bind to other locations in the γ-TuRC to propagate long-range γ-tubulin ring closure [35], but structural evidence for this hypothetical conformational state is missing.

In this study, we use cryo-electron microscopy (cryo-EM) to investigate the structure of the γ-TuRC isolated with a recombinant fragment of CDK5RAP2’s CM1 motif. We uncover a novel configuration of the γ-TuRC, in which coiled-coil CDK5RAP2 fragments are arranged into multiple tripartite assemblies together with MZT2 and GCP2-NHD subunits, which we term “CMG modules”. CMG modules dock onto the outer surface of the γ-TuRC at up to 5 binding sites, each centered on a GCP2 subunit. Remarkably, the presence of multiple CMG modules coincides with substantial displacements and rotations in γ-tubulins across the circumference of the holocomplex, wherein over half of the ∼30 nm-wide γ-TuRC adopts a conformation that closely matches that of γ-TuSC helical oligomers in the microtubule lattice-like “closed” configuration. Ring closure is achieved via the formation of novel inter-γ-tubulin and inter-GCP interfaces that are stabilized by CMG structural elements. These structural data are supported by our finding that a recombinant human γ-TuRC is potently activated by a 50 amino acid activating fragment of CDK5RAP2’s CM1 motif in a fully reconstituted system. Our results demonstrate that the GCP2 & GCP3-containing γ-TuSCs - and not just the asymmetrically arranged GCP4/5/6 subunits - are major sites of conformational regulation in the mammalian γ-TuRC, and suggest that CM1-mediated regulation of the γ-TuRC via the γ-TuSCs is evolutionarily conserved.

## Results

### The γ-TuRC can be isolated from brain tissue with a recombinant fragment of CDK5RAP2’s CM1 motif

We began by revisiting previously reported anti-γ-tubulin immunoaffinity-based purifications of γ-TuRC from mammalian brains [36,37], and asked whether γ-tubulin-containing complexes from brain tissue could be recognized by a CM1-containing polypeptide, a strategy successfully used for purifying γ-TuRC from cultured cells and *X. laevis* egg extracts [10,32,38]. Porcine brain lysates were incubated with a purified fragment of amino acids 51-100 of human CDK5RAP2 (“CDK5RAP2_51-100_”) fused to a cleavable GFP tag (“GFP-CDK5RAP2_51-100_”; [10]). A sequence alignment between *S. scrofa* and *H. sapiens* proteins indicated that CDK5RAP2_51-100_ is largely conserved between the two species, with the exception of three relatively conservative amino acid changes (T51K, H93D, and I99L, using sequence numbering from *H. sapiens*; Supplementary Figure S1A). These three residues are not resolved in the *H. sapiens* CDK5RAP2 density of a previous cryo-EM density map [12], and should therefore not disrupt the likely similar protein-protein interfaces formed between human CDK5RAP2_51-100_ and porcine γ-TuRC subunits.

Supporting this hypothesis, we successfully isolated γ-tubulin complexes from porcine brain lysates pre-incubated with GFP-CDK5RAP2_51-100_ via GFP nanobody-affinity and sucrose density gradient centrifugation (Figure 1A). The presence of high sedimentation coefficient, γ-tubulin-containing assemblies consistent with the γ-TuRC was confirmed by western blotting (Supplementary Figure S1B; [6]). Moreover, mass spectrometry (MS) of peak sucrose density gradient fractions identified the *S. scrofa* orthologs of previously identified γ-TuRC components, including γ-tubulin, GCP2, GCP3, GCP4, GCP5, GCP6, MZT1, MZT2A (hereafter, MZT2), and β-actin, as well as the exogenous human CDK5RAP2_51-100_ fragment (Supplementary Table S1). Non-erythrocyte spectrin, a previously reported component of the mammalian brain γ-TuRC [37], was also identified, albeit at a low relative abundance compared to core γ-TuRC components. In contrast to previously reported vertebrate γ-TuRCs isolated from native sources [32,39], the γ-TuRC attachment factor NEDD1 was not detected; despite this, negative stain EM revealed the presence of ∼30 nm wide ring-shaped assemblies qualitatively consistent with previous γ-TuRC structures (Supplementary Figure S1C), which is also in line with the reported dispensability of NEDD1 for the assembly of the γ-TuRC [14,16]. Lastly, we confirmed that the *S. scrofa* complexes nucleated microtubule assembly in a single-filament nucleation assay (Supplementary Figure S1D-F). Collectively, these data indicate that functional γ-TuRCs can be isolated from porcine brain tissue using CDK5RAP2_51-100_-affinity.

**Figure 1.**
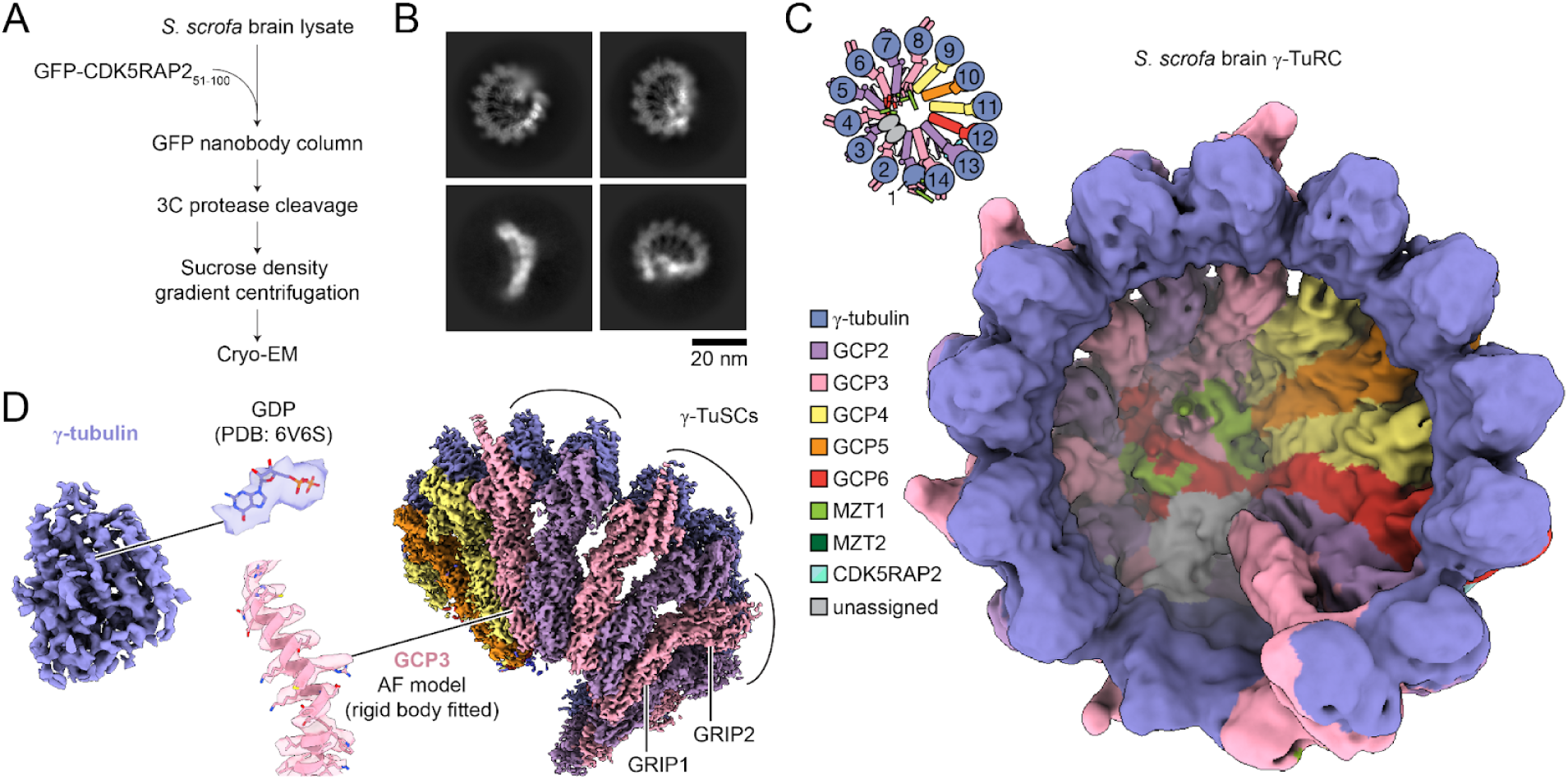
Consensus cryo-EM reconstruction of the *S. scrofa* γ-TuRC isolated from brain tissue. A) *S. scrofa* γ-TuRC isolation scheme. B) Representative 2D class averages from cryo-EM micrographs of the *S. scrofa* brain γ-TuRC. C) Overall *S. scrofa* brain γ-TuRC 3D cryo-EM reconstruction, low-pass filtered to 8 Å (surface representation). A schematic top view of γ-TuRC subunit organization is shown in the upper left corner. D) Left: Segmented γ-tubulin density (surface representation) with GDP from rigid-body fitted γ-tubulin model from PDB ID: 6V6S [10] shown in stick representation (top, middle; corresponding density for guanine nucleotide represented as transparent surface). Right: View of overall density map (surface representation) from the outside of the complex showing high-resolution features of multiple γ-tubulin and GCP subunits. Three γ-TuSC subcomplexes, as well as one of the GCP’s GRIP1 and GRIP2 domains, are indicated. A zoom in on *S. scrofa* GCP3 model predicted by AlphaFold2 and rigid-body fitted (no refinement) into the density map is shown (bottom, middle; corresponding GCP3 density represented as transparent surface) [40,41]. Densities in C)-D) are coloured according to the legend in C) and using a rigid body-docked composite model of the native human γ-TuRC [10,12].

### Cryo-EM reconstruction of the S. scrofa γ-TuRC

Next, we analyzed the structure of the *S. scrofa* γ-TuRC using cryo-electron microscopy (cryo-EM). To obtain a sufficient particle density in cryo-EM micrographs, we applied the sample multiple times onto carbon film-coated grids prior to vitrification (Supplementary Figure S2A) [10,42]. After several iterations of automated particle picking using a neural network trained on manually curated particle images followed by reference-free 2D classification (Figure 1B; [43]), we obtained a set of 456,707 final particles that led to a 4.1 Å consensus 3D reconstruction of the *S. scrofa* γ-TuRC (Figure 1C, Supplementary Figure S2B-D, and Supplementary Table S2). The consensus map has the characteristics of the γ-TuRC, including the presence of 14 densities corresponding to γ-tubulin-associated GCP subunits arranged in an asymmetric, left-handed semi-helical organization, as well as a lumenal bridge lining the inside of the cone-shaped structure. The map also contains high-resolution features, such as density for a guanine nucleotide in γ-tubulin, as well as well-defined side chains that satisfy a rigid body-docked AlphaFold prediction of *S. scrofa* GCP3 (Figure 1D; [40,41]). We note that the density for actin in the lumenal bridge is relatively weak in the overall reconstruction, indicating potential flexibility in this region of the complex. Nevertheless, the overall subunit organization of the CDK5RAP2-isolated *S. scrofa* γ-TuRC is consistent with previous reconstructions of γ-TuRCs from HeLa cell extracts [10–12], *X. laevis* egg extracts [13], as well as recombinant human complexes [14–16].

### 3D heterogeneity analysis reveals distinct subclasses of the γ-TuRC

As observed for other γ-TuRC reconstructions, the local resolution of the *S. scrofa* γ-TuRC consensus map ranges from 3 Å to >11 Å (Supplementary Figure S2B). We noticed that several regions of the map display local density features not consistent with current γ-TuRC models, particularly at positions 3, 5, and 7 of the outer face of the γ-TuRC, as well as at positions 13, 14, 1 and 2 (hereafter, the “γ-TuRC seam”), suggesting atypical structural heterogeneity in the complex. Supporting this observation, extensive 3D classification and neural network-based structural heterogeneity analyses revealed a set of three main conformations of the *S. scrofa* γ-TuRC: open, partially-assembled, and CMG-decorated (Supplementary Figure S3; [44,45]).

The first reconstruction contained the largest proportion of the consensus reconstruction particles (32%) and could be refined to a resolution of 4.3 Å (Figure 2A and Supplementary Figure S3). As expected, this class closely resembles the γ-TuRC consensus map (Figure 1C), but its resolution distribution is more uniform (Supplementary Figure S2B vs. S3). We used this set of particles to build a molecular model for the *S. scrofa* γ-TuRC (Supplementary Figure S4A and Supplementary Table S2-4; see also Figures 3 & 4). Overall, this model closely resembles the CDK5RAP2_51-100_-purified HeLa γ-TuRC [10,12], both in composition and in conformation, with the exception of subunits 13 and 14, which are in a more “sunken” conformation (Supplementary Figure S4B). This may be due to the more mobile or missing actin molecule, which could not be confidently modeled into the higher-resolution open density map due to insufficient supporting density (see Methods). Notably, the γ-tubulin ring in this subclass adopts the open configuration (Supplementary Figure S4A). We also observed recognizable, albeit low-resolution, density for the previously-assigned MZT2:GCP2-NHD bound to CDK5RAP2 at the interface of positions 12 & 13 of the γ-TuRC, into which we built a corresponding molecular model (Supplementary Figure S4). Together, these results show that this first class of *S. scrofa* γ-TuRC particles closely matches the conformation of previously described γ-TuRC reconstructions (hereafter, the “open” γ-TuRC conformation).

**Figure 2.**
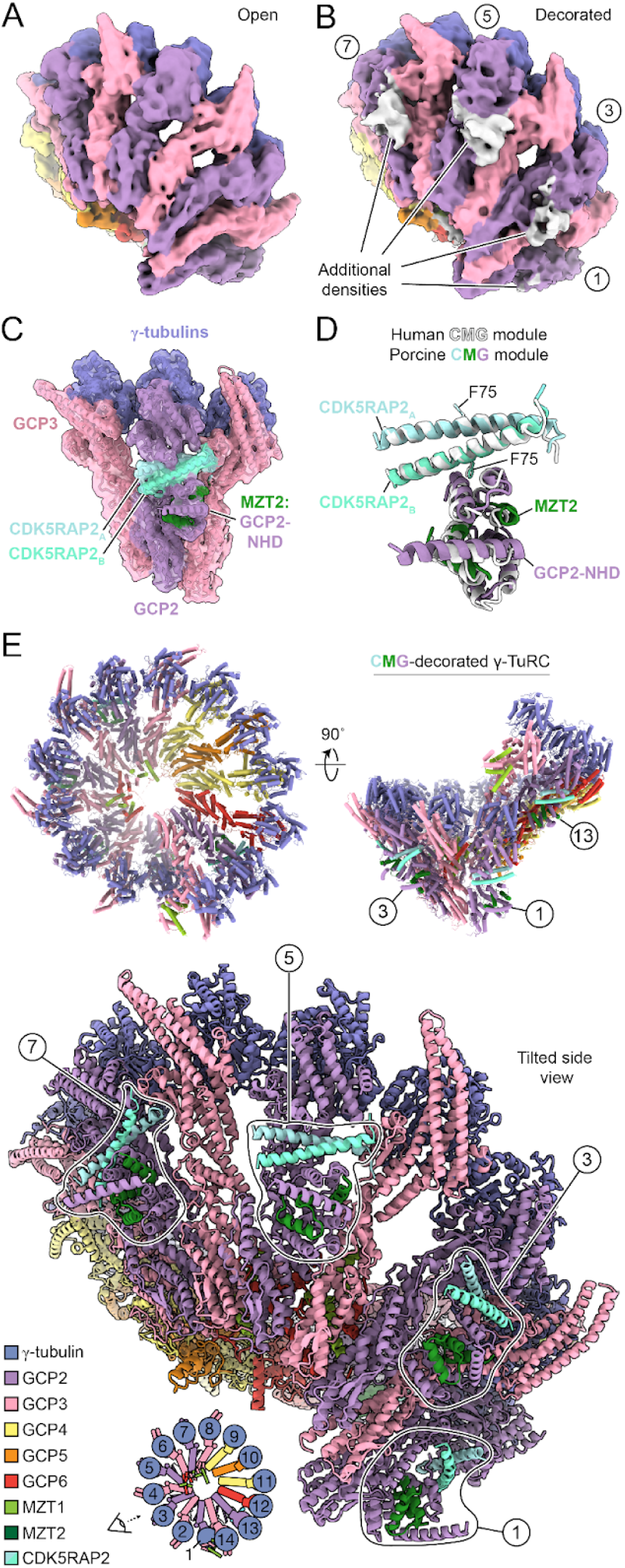
The *S. scrofa* γ-TuRC harbours multiple conformations, including a CMG-decorated state. A)-B) Cryo-EM density maps for the *S. scrofa* γ-TuRC in the “open” (A) and “CMG-decorated” conformations (B). Maps are low-pass filtered to 8 Å and viewed from a tilted angle to facilitate visualization of decorating densites. C) Transparent surface representation of a 4.9 Å focused refinement density map of CMG-decorated γ-TuRC positions 6-8, viewed from the outer face of the complex. Corresponding γ-TuRC subunit models are shown in cartoon representation. D) Cartoon representations of the *S. scrofa* CMG module at γ-TuRC position 7 aligned with the previously reported single CMG module in human γ-TuRC position 13 (coloured in white; PDB: 6X0V [12]; alignment root mean squared deviation (RMSD) = 1.3 Å). CDK5RAP2 residue F75 is shown in stick representation for both models to highlight the homology between the two models. E) Three views of a cartoon representation of the CMG-decorated *S. scrofa* γ-TuRC model. The five distinct CMG module model locations are indicated by numbers corresponding to GCP positions, as in A), and as schematized in the bottom left diagram. CMG modules at positions 1, 3, 5, and 7 are further outlined in the tilted view to emphasize corresponding locations of the additional densities in A). Colouring of subunits in A) - E) all follow the displayed legend in E).

**Figure 3.**
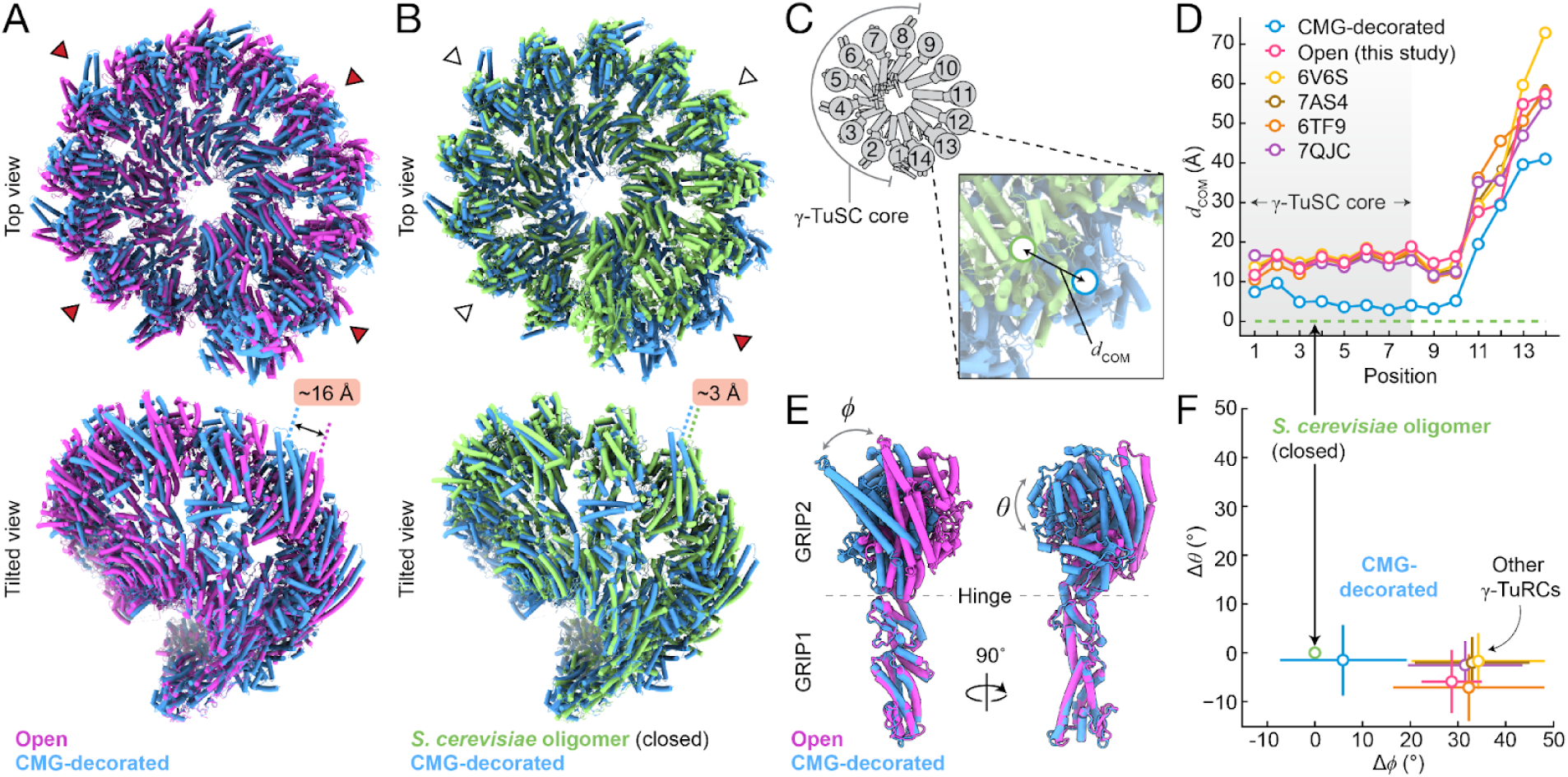
The γ-TuSC core of the CMG-decorated γ-TuRC closely resembles the structure of γ-TuSC oligomers in the closed conformation. A) Two cartoon representation views of superimposed CMG-decorated (blue) and open conformation (magenta) γ-TuRC models. B) Two cartoon representation views of superimposed CMG-decorated (blue) and closed γ-TuSC oligomer (green) pseudo-γ-TuRC models [20,29]. γ-TuRC positions with high (red triangles) and low (empty triangles) degrees of structural mismatch are indicated. Estimated Euclidian distances between γ-tubulin subunits at position 6 of the aligned complexes are also indicated A) and B). C) Schematic of the γ-TuRC showing subunit numbering. Zoom-in illustrates how Euclidian distances between γ-tubulin centers-of-mass (*d*_COM_) are used for the γ-TuRC helical parameter meta-analysis in D). γ-TuRC positions 1-8, which correspond to the γ-TuSC core, are indicated. D) Plot of *d*_COM_ versus γ-TuRC subunit position between the indicated γ-TuRC models (PDB IDs: 6V6S [10], 7AS4 [14], 6TF9 [13], 7QJC [16], as well as the open and CMG-decorated *S. scrofa* models described in this study, all relative to the closed γ-TuSC oligomer (green dashed line at *d*_COM_ = 0; [20,29]). Subunit positions 1-8 corresponding to the γ-TuSC core are indicated with the grey shaded area. E) Two cartoon representation views of superimposed models of γ-tubulin and GCP3 from γ-TuRC position 8 from the open (magenta) and CMG-decorated (blue) conformations. The radial rotation angle φ and axial rotation angle θ used to define spherical coordinates in F) are indicated. The location of the GCP hinge between N- (GRIP1) and C-terminal (GRIP2) domains is also indicated. F) Plot of the average shift in θ vs. the shift in φ for helix H12 in γ-tubulins from each γ-TuRC described in D), relative to γ-tubulins at the same positions in the closed γ-TuSC oligomer (green circle, indicated). Standard errors in φ and θ are displayed as lines. The axes in F) are scaled equally to emphasize that ring closure takes place mainly in the φ angular direction. Colouring in F) follows the legend in panel D).

The second class of *S. scrofa* γ-TuRC particles (27%) were reconstructed to an overall resolution of 4.9 Å, but the densities for GCP2, GCP3, and γ-tubulin subunits at complex positions 13, 14, 1 and 2, as well as the actin molecule, were not resolved (Supplementary Figure S5A), suggesting these subunits are highly mobile or even absent in this class. However, further subclassification revealed an additional density for a small (∼4 x ∼2 x ∼2 nm) globular domain situated on the outer face of the γ-TuRC, positioned at the vertex between GCP3 (pseudo-position 8), GCP4 (pseudo-position 9) and their associated γ-tubulins (Supplementary Figure S5A-B). The shape of this density resembles a previously described model for MZT1 in complex with the <u>N</u>-terminal α-<u>H</u>elical <u>D</u>omain of <u>GCP5</u> (GCP5-NHD) at the same location of a reconstituted human γ-TuRC assembly intermediate [16]. Using similar assignment criteria as in [16] (see Methods), we generated a locally refined density map for this region and used it to build a molecular model for *S. scrofa* MZT1 and GCP5-NHD interacting with GCP3 and GCP4 (Supplementary Figure S5C and Supplementary Tables S2-S4). The *S. scrofa* MZT1:GCP5-NHD closely resembles the human model (Supplementary Figure S5D; [16]), consistent with the reportedly high degree of structural conservation for MZT1:GCP-NHD subcomplexes [46]. In contrast with the MZT1-decorated 6-spoke structure reported by Wurtz et al. [16], we did not observe any *S. scrofa* MZT1:GCP3-NHD modules at positions 2-3, 4-5, or 6-7 accompanying our MZT1:GCP5-NHD-decorated conformation. Together with the lack of well-resolved γ-tubulin and GCP densities near the seam region, these data suggest that this particle class could represent a new γ-TuRC sub-assembly, in which a single MZT1:GCP5-NHD module decorates the outer face of the partial complex (hereafter, the “partially-assembled” reconstruction).

### Multiple CDK5RAP2 and MZT2-containing protein modules decorate the outer face of the γ-TuRC

The third reconstruction was generated from the remaining 19% of *S. scrofa* γ-TuRC particles (Figure 2B and Supplementary Figure S3). Consistent with the open and partially-assembled reconstructions, the actin density in this class is also poorly-resolved. Surprisingly, however, this class displays additional densities situated on the outer face of several GCP2 subunits (Figure 2B). Further subclassification and 3D variability analysis revealed the presence of a total of five similarly-sized densities at positions 1, 3, 5, 7, and 13 of the γ-TuRC. The density at position 7 was the most well-resolved in the initial density map, and comprises a small globular domain defined by ∼9 short α-helices, as well as an ∼8 nm-long coiled-coil, situated on the outer face of the corresponding GCP2 subunits and roughly between their N-terminal GRIP1 and C-terminal GRIP2 domain (Figure 2B). Such structural features closely resemble those for a CDK5RAP2 dimer in complex with MZT2 and the <u>N</u>-terminal α-<u>H</u>elical <u>D</u>omain of <u>GCP2</u> (GCP2-NHD) found at the interface between GCP6 and GCP2 in position 13 of a previous reconstruction of the native human γ-TuRC [12], as well as in the open conformation reported here (Supplementary Figure S4). This suggested to us that each of the additional densities observed in the third *S. scrofa* γ-TuRC particle class correspond to a similar subcomplex, but comprising exogenous human <u>C</u>DK5RAP2 associated with *S. scrofa* <u>M</u>ZT2:<u>G</u>CP2-NHD (hereafter, the “CMG module”).

To investigate this further, we generated focused refinement density maps at resolutions ranging from 4.9 – 7.6 Å targeting positions 1, 3, 5, 7, and 13 of the *S. scrofa* γ-TuRC (Supplementary Figure S3 and Supplementary Figure S6A). Supported by AlphaFold2 predictions and homology modeling (see Methods), the improved densities in the local maps were used to build and refine molecular models for five CMGs with the appropriate *S. scrofa* protein domains at γ-TuRC positions 1, 3, 5, 7, and 13, in addition to the rest of the complex (Figure 2B-C, Supplementary Figure S3 and Supplementary Table S2-4).

The *S. scrofa* γ-TuRC CMG models are similar in conformation to the previously reported human structure (RMSD = 1.3 Å between *S. scrofa* CMG at position 7 (CMG_7_) and human CMG_13_ [12]; Figure 2D). They are also similar to one another across the various CMG binding sites (e.g., RMSD = 0.8 Å between *S. scrofa* CMG_7_ vs. *S. scrofa* CMG_5_; Supplementary Figure S6A), despite differences in the surrounding interfaces formed between CMG modules and the core *S. scrofa* γ-TuRC structure (GCP3:GCP2 interface at positions 2-3, 4-5, and 6-7; no neighbouring (n-1) interface for position 1; and the GCP6:GCP2 interface at position 12-13; Figure 2D and Supplementary Figure S6A). In the CMGs, residue F75 of the GCP2 GRIP2-interfacing monomer of CDK5RAP2 (“CDK5RAP2_A_”) is correctly oriented towards a hydrophobic binding pocket formed by the C-terminal GCP2 GRIP2 domains (Figure 2D; [12]), which is essential for the CDK5RAP2:γ-TuRC interaction in cells and *in vitro* [32]. Additionally, the *S. scrofa* CMG models also now satisfy homologous, short-distance (<11.4 Å Van der Waals radius) crosslinking sites previously identified via MS between MZT2, GCP2 and GCP3 in the human γ-TuRC (e.g., between *S. scrofa* model residues MZT2 K62 & GCP2 K73; GCP2 K65 & GCP2 K347; and GCP2 K50 & GCP3 K395; [11,14]). Along with the fact that MZT2 and GCP2-NHD share a high degree of sequence identity between humans and *S. scrofa* in (>∼94% for MZT2 and >∼97% for GCP2), these observations indicate that the γ-TuRC harbours five binding sites for CDK5RAP2 and MZT2:GCP2-NHD.

### CMG module decoration is associated with a “closed” γ-TuRC conformation

Since recruitment of CDK5RAP2 to the γ-TuRC has been reported to promote the γ-TuRC’s microtubule-nucleating activity [32,35,38], we next asked whether CMG decoration influences the structure of the γ-TuRC compared to the open conformation reported here and in previous studies [10,11,13–16]. We superimposed the “CMG-decorated” *S. scrofa* γ-TuRC reconstruction with that of the open conformation using the N-terminal GRIP1 domains of GCP3 subunits at positions 2, 4, 6 and 8 as alignment anchors (see Methods). In the aligned CMG-decorated γ-TuRC, the γ-tubulins are displaced by up to ∼16 Å relative to the open conformation (Figure 3A). These γ-tubulin displacements already manifest at position 1 of the CMG-decorated complex, increase slightly through to position 9, and are sustained through the terminal γ-tubulin at position 14, as measured by comparing the change in the Euclidian distance between γ-tubulin centers-of-mass (*d*_COM_) at each position with those from the open *S. scrofa* γ-TuRC conformation, as well as with γ-tubulins from previous human γ-TuRC structures, for which PDB depositions are available (Figures 3A, C-D). The displaced γ-tubulins in the CMG-decorated complex are also accompanied by an up to ∼25° relative rotation of their associated C-terminal GCP GRIP2 domains, with the interface between GCP helices H11 and H12 (as labeled in the GCP4 crystal structure [47]) acting as a hinge point (Figure 3E). In contrast, the N-terminal GCP GRIP1 domains as well as the lumenal bridge components remain nearly identical between the CMG-decorated holocomplex and all other γ-TuRC structures (RMSD_lumenal bridge_ = 1.1 Å and RMSD_GRIP1, positions 2,4,6,8_ = 1.1 Å), emphasizing that the γ-tubulin displacements arise from the conserved hinge-like flexibility of GCPs [48].

How do the structural differences in our CMG-decorated γ-TuRC relate to the architecture of the canonical 13-protofilament microtubule architecture found in most cell types [49]? To address this question, we built a pseudo-γ-TuRC model from repeating *S. cerevisiae* γ-TuSC subunits in the “closed” conformation, which matches closely with the helical geometry of a 13-protofilament microtubule [20,29], and included this model in our N-terminal GCP3 GRIP1 domain-aligned γ-TuRC meta-analysis. This revealed that the γ-tubulins in the CMG-decorated γ-TuRC are overall much closer to the microtubule geometry than all other vertebrate γ-TuRC structures, including the open conformation reported here (average relative displacement = 5.0 ± 2.0 Å for CMG-decorated vs. 14.9 ± 2.1 Å for the other γ-TuRCs; Figure 3C-D). We also calculated the distribution of angular rotations in each γ-tubulin subunit relative to its analog in the closed pseudo-γ-TuRC model (see Methods), and found that the CMG-decorated γ-tubulins are closely aligned (Figure 3E-F). In contrast, the average change in the angular rotation of γ-tubulins in the other structures, including the S. scrofa open γ-TuRC, all cluster farther away from the closed pseudo-γ-TuRC (average radial angular displacement (Δφ) = 6.0° for CMG-decorated vs. 31.9° for the other γ-TuRCs; average axial angular displacement (Δθ) = -1.5° for CMG-decorated vs. -3.2° for the other γ-TuRCs; Figure 3F). Thus, our analysis shows that the CMG-decorated γ-TuRC undergoes a substantial set of conformational changes, in which the majority of its γ-tubulin subunits are reorganized into a configuration that closely matches the architecture of a 13-protofilament microtubule.

### MZT2:GCP2-NHD together with CDK5RAP2 form a molecular wedge that stabilizes γ-tubulin ring closure

To understand how CMG module decoration correlates with γ-tubulin ring closure, we next compared γ-TuRC subunit interfaces in the open and CMG-decorated conformations. Several notable differences were observed. First, the interface formed between γ-tubulins is altered in the CMG-decorated γ-TuRC. In the open conformation, this interface is mainly formed by interactions between α-helices H6 and H9 of one γ-tubulin with α-helices H2’’ and H3 of the neighbouring γ-tubulin [10], but in the CMG-decorated complex, the neighbouring γ-tubulin is vertically displaced by up to 17 Å (as judged by the distance between Cα of Arg343 (Supplementary Figure S6B-C). This shift places H2’’ and H3 near the vicinity of the bottom parts of H6 and H9, which is also accommodated by structural rearrangements in the neighbouring GCPs (Figure 3E and S6B-C). Accordingly, the staggered positioning between γ-tubulins apparent in all open γ-TuRC structures is smoothed out in the CMG-decorated complex (mean variance in neighbour-to-neighbour displacement = 11 Å for open vs. 4 Å for CMG-decorated conformations; Figure 3D). These observations reveal a high degree of structural plasticity in the lateral interfaces formed between γ-tubulin neighbours in the vertebrate γ-TuRC, a property also previously described for interfaces between α- and β-tubulin in the microtubule [17].

Second, the N-terminal ends of both copies (A and B) of CDK5RAP2 within the CMG module form contacts with a conserved structural feature - which we call the GCP “spoon” - in the neighbouring (position n-1) GCP subunit. In the case of positions 7, 5, and 3, this feature is presented by GCP3 (Figure 4A-B), while for position 13, the GCP6 spoon is presented. In all the GCPs, the spoon consists of two short, antiparallel α-helices connected by a semi-structured loop that, in the open conformation, contacts both the neighbouring (position n+1) GCP subunit and the minus end of its associated γ-tubulin (e.g., Figure 4B & E). The spoon also contains 2 antiparallel β-strands, which face the CDK5RAP2 dimer binding pocket of the (n+1)^th^ GCP subunit (Figure 4B and Supplementary Figure S6D), so it is well-positioned to stabilize CDK5RAP2-induced structural reorganization of GCP2’s GRIP2 domain.

**Figure 4.**
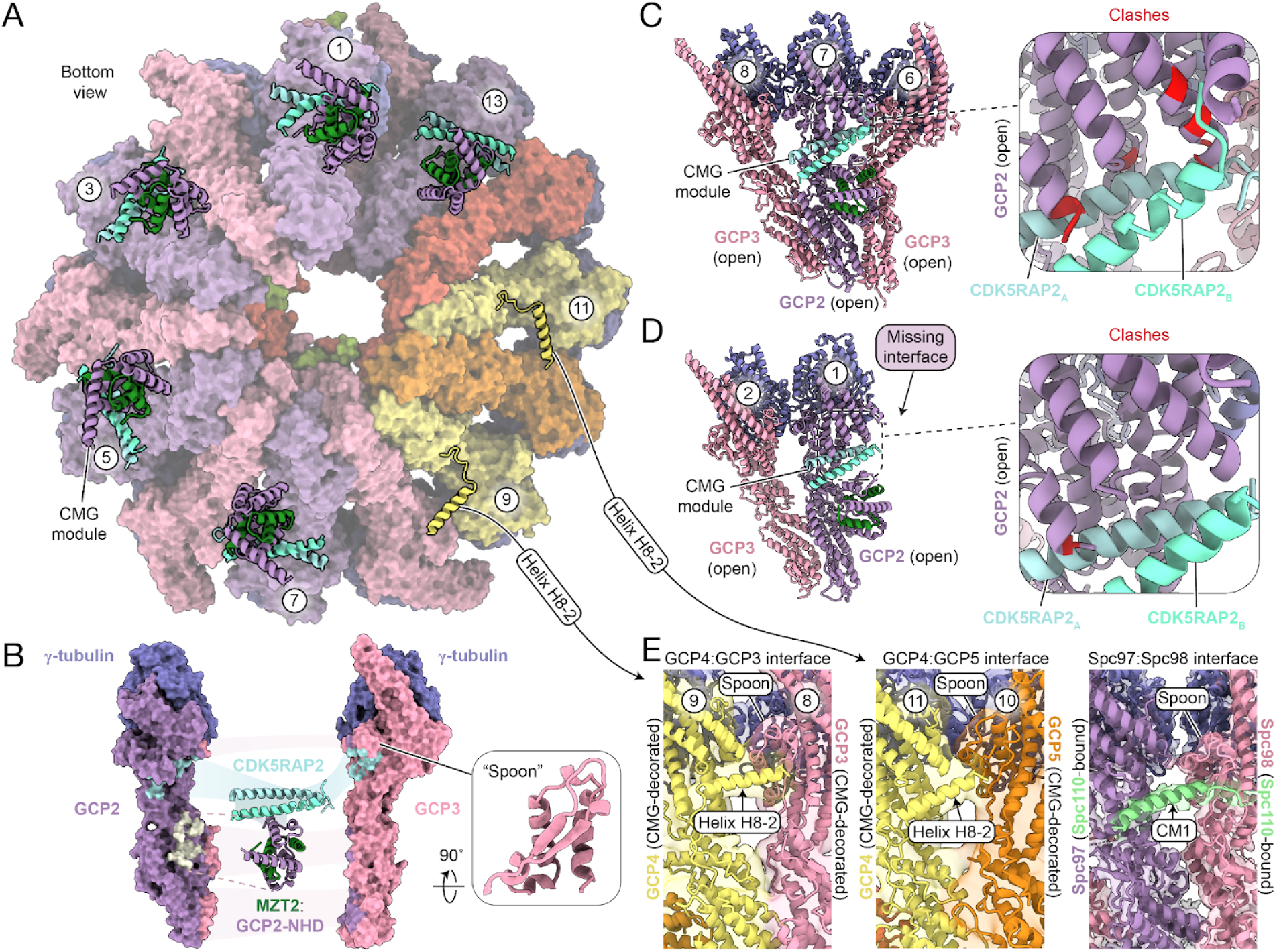
The CMG module forms a molecular wedge that promotes γ-tubulin ring closure. A) Faded surface representation of the CMG-decorated γ-TuRC viewed from the bottom of the complex to facilitate visualization of GCP binding sites on the outer face of the holocomplex. CMG modules at positions 1, 3, 5, 7, and 13, as well as the structure helix H8-2 of both GCP4 subunits at positions 9 and 11 are indicated and shown as overlaid cartoon representations. B) Exploded views of the CMG (cartoon representation)-decorated γ-TuSC (surface representation) at positions 7-8. Subunits are coloured as in Figure 2, except for interfaces, which are highlighted and coloured according to the interacting subunit. Inset shows a cartoon representation, rotated view of the GCP3 “spoon” to highlight secondary structure features. C) Cartoon representation overview of GCP2, GCP3 and γ-tubulins from positions 6-8 (indicated) of the open γ-TuRC conformation and viewed from the outer face of the complex. The CMG module from the same subunits in the aligned CMG-decorated γ-TuRC is also displayed and indicated. Right: Zoomed in view of the CMG module showing clashing residues (within <0.6 A of calculated Van der Waals radii; labeled and coloured in red) in the open conformation GCP2 subunit. D) Cartoon representation overview of GCP2, GCP3 and γ-tubulins from positions 1-2 (indicated) of the open γ-TuRC conformation and viewed from the outer face of the complex. The CMG module from the same subunits in the aligned CMG-decorated γ-TuRC is also displayed and indicated. Right: Zoomed in view of the CMG module showing clashing residues on GCP2 as in C). E) Left: Cartoon representation of the GCP4:GCP3 interface from positions 8-9 of the CMG-decorated γ-TuRC in the corresponding cryo-EM density (transparent surfaces). Arrow indicates GCP4 helix H8-2. Middle: Cartoon representation of the GCP4:GCP5 interface from positions 10-11 of the CMG-decorated γ-TuRC in the cryo-EM density (transparent surfaces). Arrow indicates GCP4 helix H8-2. Right: Cartoon representation of the reconstructed Spc97:Spc98 interface from the *S. cerevisiae* γ-TuSC oligomer in the corresponding cryo-EM density (transparent surfaces; PDB ID: 7M2Y [28]). Arrow indicates the Spc110 CM1 motif. The GCP spoon feature is indicated in all three interfaces.

Third, when comparing the structure of GCP2 in position 7 of the open and CMG-decorated γ-TuRCs, the C-terminal GRIP2 domain is shifted up and rotated to accommodate CMG module-binding (Figures 3A & E); otherwise, CDK5RAP2_A_ would clash with multiple α-helices in GCP2 (Figure 4C). Unexpectedly, far fewer clashes are observed between the CMG module and GCP2 in the open conformation at position 1 (Figure 4D), and the γ-tubulin:γ-tubulin interface at positions 1:2 is similar to the one in the open γ-TuRC conformation. We note that this position misses a position (n-1) GCP binding partner (and hence, a GCP spoon) - for CDK5RAP2 to interact with (Figure 4D), further emphasizing the potential role of the GCP spoon in CMG module function.

Fourth, we focused on the interface between CDK5RAP2_B_ and MZT2:GCP2-NHD within the CMG module. We attempted to reconstitute the CMG module with recombinant truncation constructs in GST pulldown experiments. We expressed and purified a fragment of CDK5RAP2 encompassing the resolved portion of its α-helical coiled-coil in the CMG module (CDK5RAP2_44-93_), as well as MZT2 in complex with GST-GCP2-NHD, from bacteria (Supplementary Figure S7A-B). Surprisingly, GST-tagged MZT2:GCP2-NHD was not able to pull down detectable quantities of CDK5RAP2_44-93_, which, considering the concentrations tested and the detection limit of Coomassie staining, suggests an affinity of at best ∼10 μM (Supplementary Figure S7C). We note that, compared with the GCP2 GRIP1 and GRIP2 domains, GCP2-NHD is less well-conserved, including at residues 1-35, which form the CDK5RAP2**_B_**-binding interface (Supplementary Figure S7D-F). From our structural models, the calculated binding energy of this particular interface is ∼1.6 kcal mol^-1^ lower than, for example, that of the interface formed between CDK5RAP2_A_ and the GCP2 GRIP2 domain (-8.3 kcal mol^-1^ for CDK5RAP2_A_:GCP2 interface, vs. -6.7 kcal mol^-1^ for CDK5RAP2_B_:GCP2-NHD interface, which at 4° C correspond affinities of ∼0.26 μM vs. ∼4.8 μM, respectively; calculations use CMG_7_ [50]). Altogether, these observations lead us to propose that MZT2:GCP2-NHD forms a loosely defined interface with CDK5RAP2_B_, which requires additional interactions between CDK5RAP2_A_, the GCP2 GRIP2 domain, and the (n-1)^th^ GCP spoon to promote γ-tubulin ring closure in the CMG-decorated γ-TuRC.

Lastly, when generating initial AlphaFold2 models for *S. scrofa* GCP4 (see Methods), we noticed that an α-helix between β-strands β3 and β4 predicted for both the human and *S. scrofa* GCP4 (residues 228-250; hereafter, “helix H8-2”) is not resolved in the open conformation density map (Supplementary Figure S8A-B). Helix H8-2 is also missing in the human GCP4 crystal structure [47], as well as in all currently available vertebrate γ-TuRC cryo-EM reconstructions [10,13,14,16]. In contrast, we observed clear density for helix H8-2 in both copies of GCP4 (positions 9 and 11) in the CMG-decorated γ-TuRC (Figure 4A & E). Surprisingly, this α-helix occupies the binding site of CDK5RAP2 in GCP2, as well as the binding site of *S. cerevisiae* Spc110’s CM1 motif in Spc97 (Figure 4E) [28]. The conformation of helix H8-2 is more similar to Spc110’s CM1 motif than to CDK5RAP2_51-100_ (Supplementary Figure S8C & D). Analogous to CDK5RAP2 and Spc110, both copies of H8-2 also contact the spoon elements in the neighbouring GCP subunits (GCP3 for H8-2 at position 9, and GCP5 for H8-2 at position 11; Figure 4E and Supplementary Figure S6D). Although the sequence conservation between helix H8-2 and the CM1 motifs is low (<∼20% identity), the α-helix itself is conserved in GCP4s from multiple species (Supplementary Figure S8E). These findings show that GCP4 helix H8-2 becomes structured in the CMG-decorated γ-TuRC, raising the possibility that certain GCPs may have evolved structural mimicry to perform CM1-like functions via internal motifs in the γ-TuRC.

### Untagged CDK5RAP2 promotes microtubule nucleation from both the native S. scrofa complex and a fully reconstituted human γ-TuRC

The “closed” conformation of γ-TuSC subunits in the CMG-decorated *S. scrofa* γ-TuRC more closely resembles the 13-protofilament microtubule lattice [17], and this conformation is stabilized in part by CDK5RAP2. We reasoned that since only a fraction (∼20%) of the *S. scrofa* γ-TuRC particles are found in a CMG-decorated state by cryo-EM, the majority of γ-TuRCs could still transition into this conformation, even if transiently, and we could sample these additional activation events in a single filament assay. We therefore next tested the activity of the *S. scrofa* γ-TuRC non-specifically adsorbed to the silanized glass and introduced soluble fluorescently-labeled tubulin and GTP to initiate microtubule polymerization, as in our initial characterization of the complex (Supplementary Figure S1D), but in the presence of CDK5RAP2_44-93_ (Supplementary Figure S7A). Importantly, this CDK5RAP2 construct contains neither bulky nor charged affinity tags after purification, both of which have reportedly inconsistent effects on γ-TuRC binding and activation [13,21,38].

Gratifyingly, in the presence of 2 μM CDK5RAP2_44-93_, the *S. scrofa* γ-TuRC nucleated microtubules at levels which were several orders of magnitude higher than in controls (13 ± 9 microtubules per field of view in the presence of 2 μM CDK5RAP2_44-93_ vs. 0 ± 0 microtubules per field of view in the control; Supplementary Figure S9A). To ensure that activation was specific to CDK5RAP2_44-93_, we also tested a construct containing an F75A mutation (Supplementary Figure S9B), which had previously been shown to disrupt γ-TuRC-binding and activation [38], and found that it did not promote microtubule nucleation from the *S. scrofa* γ-TuRC (Supplementary Figure S9A).

The *S. scrofa* γ-TuRC was affinity-isolated from brain tissue using CDK5RAP2_51-100_, which already contains some fraction of activator-bound complexes (Figure 2 and Supplementary Figure S3). To test whether activation is a general feature of the γ-TuRC, we next examined the activation potential of human γ-TuRC reconstituted in insect cells [15]. We purified recombinant γ-TuRC (rec-γ-TuRC) reconstituted with a defined set of human components (Supplementary Figure S9C), including MZT2 (see Methods), and confirmed that it lacks any insect-derived proteins by MS, as previously reported [15]. We immobilized rec-γ-TuRC to passivated coverslip surfaces via a monomeric EGFP tag fused to its MZT2 subunits and acquired two-colour time lapse images of the microtubule polymerization reactions (Figure 5A and Supplementary Figure S9D). In control experiments, the percentage of rec-γ-TuRC GFP puncta that nucleated microtubule assembly was relatively low (0.02 ± 0.02 microtubules per GFP punctum; Figure 5A & E and Supplementary Figure S9D), consistent with previous reports that the γ-TuRC is a poor nucleator in the absence of activating factors [11,15,18,19,21,32,35,38].

**Figure 5.**
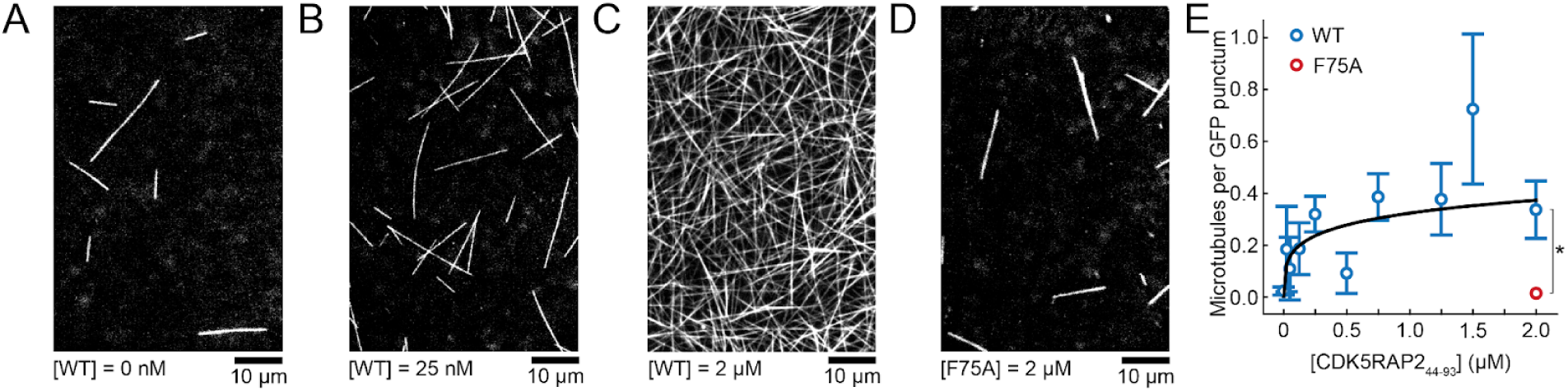
CDK5RAP2_44-93_ promotes microtubule nucleation from recombinant human γ-TuRC. A) - D) Fields of view from TIRF microscopy experiments, in which dynamic microtubules are nucleated by surface-immobilized rec-γ-TuRC. The images show the fluorescent tubulin channel at t = 21 min in the presence of no added wild-type (WT) CDK5RAP2_44-93_ (A), 25 nM WT CDK5RAP2_44-93_ (B), 2 μM WT CDK5RAP2_44-93_ (C), or 2 μM of the CDK5RAP2_44-93_ F75A mutant (“F75A”) (D). E) Plot of the number of microtubules divided by the number of detected γ-TuRC-GFP puncta at t = 21 min in the presence of increasing amounts of WT CDK5RAP2_44-93_ (blue circles). The solid line is a fit to the Hill equation. Data for the F75A mutation are shown in red, which were statistically significantly different from the WT data at the same CDK5RAP2_44-93_ concentration (single asterisk: p-value = <0.000001 from a standard t-test), but not from the WT data with no added CDK5RAP2_44-93_ (p-value = 0.32 from a standard t-test). Each data point represents the mean ± standard error of between 6-9 fields of view sampled from between 2-3 independent experiments. CDK5RAP2 concentrations are reported for the dimer, and the tubulin concentration in all experiments was 12.5 μM.

In the presence of increasing amounts of CDK5RAP2_44-93_, we observed a dose-dependent increase in the number of microtubules nucleated by rec-γ-TuRC (0.3 ± 0.1 microtubules per GFP punctum in the presence of 2 μM CDK5RAP2_44-93_; [CDK5RAP2_44-93_] at half-maximal activation = ∼100 nM; Figure 5B-C & E). The F75A mutant did not promote microtubule nucleation from rec-γ-TuRC (0.02 ± 0.01 microtubules per GFP punctum in the presence of 2 μM F75A-mutated CDK5RAP2_44-93_; Figure 5E). As a further control, we confirmed that CDK5RAP2_44-93_ itself did not promote microtubule nucleation in the absence of *S. scrofa* γ-TuRC or rec-γ-TuRC (no microtubules observed in 9 fields of view from 3 independent experiments). Together, these data clearly demonstrate that CDK5RAP2_44-93_ potently activates the microtubule-nucleating activity of: (i) the *S. scrofa* γ-TuRC; and (ii) a human γ-TuRC in a fully reconstituted system and at concentrations comparable to full-length CDK5RAP2 [35].

## Discussion

In this study, we successfully isolated the native γ-TuRC from porcine brains with a fragment of CDK5RAP2’s CM1 motif and determined its structure using cryo-EM. In addition to validating mammalian brain tissue as an attractive source of native γ-TuRC material, our structural data have led to a model for the CMG-decorated conformation of the complex distinguished by a more closed γ-tubulin ring than in previously-determined γ-TuRC reconstructions (Supplementary Video S1). This new conformation provides structural evidence that accessory proteins can transition the γ-TuRC into a closed state in the absence of α/β-tubulin [35], and provides an important view into the activation pathway of the γ-TuRC, which has consistently exhibited surprisingly low basal microtubule-nucleating activity *in vitro*.

Previous work has suggested that the γ-TuSC core of the γ-TuRC (positions 1-8) is structurally more well-suited to nucleate a microtubule, and that the asymmetrically-positioned GCP4/5/6 γ-TuRC subunits could provide binding sites for regulatory factors [10,11,15]. Our meta-analysis of available γ-TuRC structures shows that all previous γ-TuRC structures exhibit an open γ-tubulin ring conformation, even in the γ-TuSC core, which far from matches the arrangement of α/β-tubulin in the 13-protofilament microtubule lattice (Figure 3). The CMG-decorated conformation most closely matches the microtubule lattice at positions that encompass the γ-TuSC core (i.e., positions ∼1-10). While our data do not rule out the possibility of further regulation at other sites in the complex, our work does reveal that the evolutionarily conserved γ-TuSCs are sites of major conformational regulation in the vertebrate γ-TuRC, potentially explaining why the loss of GCP4/5/6 - but not CM1 motif proteins - has been tolerated in certain fungal lineages [9].

We note that the remaining γ-tubulins in the CMG-decorated complex (positions 11-14) still deviate from the microtubule architecture, raising the question of whether and/or how these positions undergo ring closure. GTP-binding in γ-tubulin is critical for microtubule nucleation and may play a role [51,52], although the potential mechanisms are still unclear. Strong cooperativity in basal microtubule nucleation rates in response to increasing α/β-tubulin levels indicates that the presence of substrate could also help bypass structural and kinetic hurdles [11,15,53]. This is almost certain to be a key additional transition step, as individual γ-tubulins in our CMG-decorated complex still exhibit secondary structure features reminiscent of unpolymerized β-tubulin (Supplementary Figure S6B-C; [54,55]), which in a recent preprint has been proposed to be corrected after the γ-TuRC nucleates the microtubule and remains bound as a minus end cap [56]. An intriguing result from our CMG-decorated reconstruction is the structuring of GCP4 helix H8-2, which occupies the CM1 binding pocket of both copies of GCP4 and contacts the spoon element of the neighbouring GCP subunit at the (n-1)^th^ positions (Figure 4). Post-translational modifications and/or the formation of analogous interfaces with unidentified binding partner(s) as seen in the CMG module may help induce γ-tubulin ring closure at these sites; i.e., every odd-positioned site in the complex (1, 3, 5, 7, 9, 11 and 13) may exhibit a “ring-closing” feature.

Our finding that the CMG-decorated γ-TuRC is decorated with five sub-assemblies containing CDK5RAP2, MZT2, and GCP2-NHD helps provide the most complete molecular picture to date of how CM1 motifs elicit their function in the vertebrate γ-TuRC. In addition to these structural data, we provide additional observations: 1) CDK5RAP2_44-93_ activates both the fractionally CMG-occupied *S. scrofa* γ-TuRC material and, at nM concentrations, the reconstituted human γ-TuRC (Figure 5E and Supplementary Figure S9D); and 2) CDK5RAP2_44-93_ does not appreciably interact with a reconstituted MZT2:GCP2-NHD.

This suggests that the CMG module as a whole may be a transient structure, which would be consistent with difficulties in observing the fully-decorated structural state by cryo-EM both here and in previous work [10,12,13]. Our findings suggest a model, in which energetically more-favourable interfaces formed between CDK5RAP2, GCP2’s C-terminal GRIP2 domain, and the neighbouring GCP spoon, are transiently stabilized by the presence of MZT2:GCP2-NHD, all of which come together to form a molecular wedge that can prevent the transition of GCPs back to the open conformation. Given that templated nucleation is a kinetically unfavourable yet highly cooperative process [11,15,23,53], transient and short-lived CMG-decorated γ-TuRC conformations may be sufficient to achieve MTOC-localized microtubule nucleating activity, while still offering a way to suppress nucleation in cytosol.

Previous work using a similar CDK5RAP2-based purification strategy identified only a single CMG module at position 13 of the HeLa complex [10]. Similarly, the *S. scrofa* open conformation still has a CMG module at the same position (Supplementary Figure S4). Together with our structural data, our finding that the microtubule-nucleating activity of the brain γ-TuRC can be enhanced by addition of CDK5RAP2_44-93_ suggests that this occurs via CMG module formation and docking at positions 3, 5, and 7. Thus, the position 13 CMG module may not be as important for activating the γ-TuRC’s microtubule-nucleating function as other CMG sites. The CMG module at position 13 is also unique, because it interacts with the GCP6 spoon instead of the GCP3 spoon. This site could instead be important for CDK5RAP2’s γ-TuRC-tethering function at the centrosome, facilitated by e.g. conserved CDK5RAP2-pericentrin interactions [33], as well as phosphoregulation of CDK5RAP2 autoinhibition [34,57]. Once the γ-TuRC is recruited, a high local concentration of centrosomal CDK5RAP2 and other regulators, which may arise from biomolecular condensation of its low-complexity domains [58], could shift the equilibrium towards CMG module formation at other sites on the γ-TuRC followed by γ-tubulin ring closure and activation. This model helps explain why overexpression of CDK5RAP2 fragments results in dramatic ectopic microtubule nucleation in the cytosol [32]; i.e., inactive cytoplasmic complexes become artificially exposed to a centrosome-like environment.

Our work highlights the role of long-range allostery in regulating γ-TuRC activity. Allosteric activation of a nucleation template by nucleation-promoting factors (NPFs) is also observed for the actin filament-nucleating Arp2/3 complex. For example, several members of the WASP family of proteins have been shown to promote a conformational change that repositions its actin-like subunits into a short-pitch conformation resembling the actin filament [59]. However, complete activation requires additional tethering of incoming actin [60]. Interestingly, full-length CDK5RAP2 has been shown to not only stimulate microtubule nucleation from the γ-TuRC, but also promotes its ability to cap the microtubule minus end [35]. Our work spurs the question of whether oligomeric binding of CDK5RAP2 and/or other γ-TuRC-associated proteins could perform similar α/β-tubulin-recruitment and tethering functions at MTOCs. A recent preprint describing a subtomogram-averaged structure of *S. cerevisiae* spindle pole microtubules capped by the γ-TuRC demonstrated the presence of a coiled-coil protein linking alternating γ-TuRC subunits with α/β-tubulin at the minus end [61], which we note is reminiscent of the odd-numbered subunit CM1 motif-like binding in our CMG-decorated model (Figure 4A). Further structural investigations of *in vitro*-reconstituted as well as *in situ* microtubule end-capped γ-TuRC co-complex structures will undoubtedly shed light onto the regulation of microtubule nucleation at diverse MTOCs.

## Acknowledgements

We thank Daniel Böhringer and Pavel Afanasyev from the Cryo-EM Knowledge Hub (ETH Zürich) for cryo-EM image processing advice, as well as Miroslav Peterek and Bilal Qureshi from ScopeM (ETH Zürich) for cryo-EM sample preparation and data collection training and support. We also thank Dorothea Pinotsi (ETH Zürich) from ScopeM for fluorescence microscopy training and support. We are grateful to Stadt Zürich, Umwelt- und Gesundheitsschutz, Fachbereich Veterinärdienste for supplying porcine brain tissue. We thank Aitor Pellicer Camardiel, Mirko Markovic, Bin Tsai, and members of the Pilhofer Group at ETH Zürich for helpful discussions, Stefanie Jonas for critical reading of the manuscript, as well as Christian Landolt and Alex Myczko (ETH Zürich) for IT support. This study includes calculations performed on the Euler cluster of ETH Zürich. Proteomics experiments were performed at the Functional Genomics Center Zurich (FGCZ) of University of Zürich and ETH Zürich. This work was supported by startup funds from the ETH Zürich, an SNSF Project Grant (#310030_208120), and an SNSF Starting Grant (#TMSGI3_211309), awarded to MW. FC was financially supported by the ETH Zürich Amgen Scholars program funded by the Amgen Foundation.

## Author Contributions

YX, RK and MW designed the study, and purified & characterized all proteins used. YX and HMH collected cryo-EM data. HMH and MW performed cryo-EM processing. HMH, FC, and RK built structural models. YX, RK and MW designed and cloned protein constructs. YX, RK, FM, and MW performed TIRF microscopy experiments. FM wrote and optimized the helical analysis and TIRF microscopy analysis python scripts. YX and MW wrote the paper with contributions from other authors. MW supervised the study and obtained funding.

## Materials and Methods

### Isolation of *S. scrofa* brain γ-TuRC

Porcine brains were obtained from a local slaughterhouse (Stadt Zürich, Umwelt- und Gesundheitsschutz, Fachbereich Veterinärdienste) and were processed for long term storage following the method described previously for porcine brain dynactin complex isolation [62]. Briefly, fresh brains were placed into ice cold 1X PBS less than 2 hr after slaughter. Blood vessels and meninges were removed and brains were washed in 1X PBS followed by HB buffer (35 mM PIPES, pH 7.2, 5 mM MgSO4, 100 µM EGTA, 50 µM EDTA). Brains were then individually snap frozen in liquid nitrogen and stored at -80 °C.

The isolation method of native γ-TuRCs from porcine brains followed the previously described strategy for human γ-TuRCs isolated from HeLa cell cytoplasmic extracts [10]. Briefly, a recombinant fragment comprising part of the CM1 motif of human CDK5RAP2_51-100_ (also referred to as γ-TuNA [32]) fused to a cleavable GFP tag (His6-SUMO-TEV-GFP-PreScission-γ-TuNA, a gift from Tarun Kapoor (Addgene plasmid # 178066 ; http://n2t.net/addgene:178066 ; RRID:Addgene_178066)) was transformed into BL21-CodonPlus (DE3)-RIL (Stratagene) cells, expressed in 6 L of LB broth and purified using Ni-NTA affinity, TEV protease cleave, and size exclusion chromatography, identically as described [10]. An N8his-GFPenhancer-GGGGS4-LaG16 plasmid for production of a tandem GFP nanobody was a gift from Motoyuki Hattori (Addgene plasmid # 140442 ; http://n2t.net/addgene:140442 ; RRID:Addgene_140442). The plasmid was transformed into BL21-CodonPlus (DE3)-RIL (Stratagene) cells and expressed in 12 L of LB after induction with 0.5 mM IPTG at 18 °C for 16 hr. The nanobody purification method was modified slightly from [10]. Cells were first harvested by centrifugation at 4000 g at 4 °C for 10 min. The pellet was resuspended in lysis buffer (50 mM Tris pH 8.0, 150 mM NaCl) containing 1 mM PMSF and lysed by passing through an Emulsiflex C5 microfuildizer (Avestin) 3 times at >10,000 psi. The lysate was centrifuged at 45,000 rpm in a Type 45 Ti rotor (Beckman) for 1 hr at 4°C. The supernatant was incubated with 3 mL Ni-NTA agarose (Qiagen) pre-equilibrated with binding buffer (50 mM Tris-HCl pH 8.0, 150 mM NaCl, 30 mM imidazole) via nutation at 4 °C for 1 hr. The resin was then washed with 10 CV of binding buffer and the nanobody was eluted using an elution buffer (50 mM Tris-HCl pH 8.0, 150 mM NaCl, 300 mM imidazole). The eluate was pooled and digested with 200 µg of HRV 3C protease produced in-house while dialyzing into dialysis buffer (50 mM Tris-HCl pH 8.0, 150 mM NaCl, 15 mM imidazole) in a 3.5 kDa cut-off dialysis membrane overnight at 4 °C. The digested protein was then manually loaded onto a 1 mL HisTrap HP column (Cytiva) pre-equilibrated with dialysis buffer using a syringe. Protein-rich fractions were identified via Bradford assay [63], and concentrated to <1 mL using an Amicon Ultra 10 kDa cut-off centrifugal filter. Concentrated protein was applied to a Superdex 75 Increase 10/300 size-exclusion column (Cytiva) pre-equilibrated with gel filtration buffer (100 mM NaHCO_3_ pH 8.0, 150 mM NaCl). Peak fractions were pooled and concentrated to 10 mg/ml. 10 mg of purified GFP nanobody was then coupled to a 1 mL NHS HiTrap HP column (Cytiva) following the manufacturer’s instructions and stored in storage buffer (1X PBS, 1 mM NaN_3_) at 4 °C.

Frozen porcine brains (typically between 1-3) were thawed on ice and blended in 100 mL lysis buffer (50 mM HEPES, pH 7.5, 150 mM NaCl, 1 mM MgCl2, 1 mM EGTA, 0.1% IGEPAL, 0.1 mM GTP) per brain. The lysate was centrifuged in a JLA-16.250 rotor (Beckman) at 16,000 rpm for 15 min at 4 °C. The supernatant was collected and centrifuged in a Type 45 Ti rotor (Beckman) at 45,000 rpm for 50 min at 4 °C. Clarified lysate was mixed with 10 mg purified GFP-γ-TuNA and incubated with nutation for 1 hr at 4 °C. The mixture was filtered over a 0.2 µm PVDF syringe filter and loaded onto the GFP nanobody column pre-equilibrated with lysis buffer. The column was then washed with 50 mL of lysis buffer. 100 ug HRV 3C protease was diluted into 1 mL of lysis buffer and subsequently injected into the column. Proteolytic digestion of the GFP tag to release complexes was allowed to proceed overnight at 4 °C. γ-TuRCs were eluted in 100 uL fractions with lysis buffer. The 3 peak fractions identified based on Bradford assay were pooled and layered onto a 2 mL discontinuous sucrose gradient containing 10%, 20%, 30%, and 40% (w/v) sucrose in gradient buffer (40 mM HEPES, pH 7.5, 150 mM NaCl, 1 mM MgCl2, 1 mM EGTA, 2 mM 2-mercaptoethanol, 0.01% IGEPAL, 0.1 mM GTP) layered in TLS-55 (Beckman) centrifuge tubes. The gradient was centrifuged in a TLS-55 rotor at 50,000 rpm for 3 hr at 4°C with no brake and carefully fractionated into 250 μL fractions from the top of a gradient with a cut-off P1000 pipette tip. Fractions were analyzed by SDS-PAGE followed by Coomassie and/or silver-staining (Thermo Fisher, catalog number:24612) and via anti-γ-tubulin western blot. Peak fractions (typically fraction 6 & 7 from the top of the gradient; Supplemental Figure 1B) were aliquoted and snap frozen in liquid nitrogen and stored at -80°C. For estimating sedimentation coefficients, parallel gradients were run with 200 µg each of BSA, catalase, and thyroglobulin.

### Purification of recombinant, untagged CDK5RAP2 fragments

A bacterial expression construct for amino acids 44-93 of CDK5RAP2 lacking a GFP tag was generated by first PCR amplifying CDK5RAP2_44-93_ from pRcCMV Cep215 (NCBI accession BAB13459.3), a gift from Erich Nigg (Addgene plasmid # 41152; http://n2t.net/addgene:41152; RRID:Addgene_41152) using the following primers: 3’-CGTAGTCGACCGGTTCTTCTCTCCCAAATGTGTCAGAAGA-5’ and 3’-CGTAGCGGCCGCTCAATGAAATTCCTGTTGCATTC-5’. The PCR product was inserted into the SalI and NotI sites of a modified pET-SUMO vector (pET His6 Sumo TEV LIC cloning vector (1S), a gift from Scott Gradia (Addgene plasmid # 29659; http://n2t.net/addgene:29659; RRID:Addgene_29659) using standard restriction cloning techniques, generating a construct for expressing His6-SUMO-TEV-CDK5RAP2_44-93_ in bacteria. The plasmid was sequence verified using Sanger sequencing (Microsynth).

The plasmid was transformed into BL21(DE3) pRIL *E. coli* cells (Stratagene) and His6-SUMO-TEV-CDK5RAP2_44-93_ expression was induced with 0.5 mM IPTG for 16 hours at 18°C. Cell pellets from 3 L culture were resuspended in 45 mL Ni-NTA lysis buffer (50 mM sodium phosphate, pH 8.0, 300 mM NaCl, 15 mM imidazole, 0.1% (v/v) Tween-20 and 1 mM 2-mercaptoethanol) and lysed by three passes through an Emulsiflex C-5 (Avestin). Lysate was clarified at 35,000 rpm in a Type 45 Ti rotor (Beckman) for 45 min at 4°C and the supernatant was mixed with 3 mL Ni-NTA agarose (Qiagen) pre-equilibrated in lysis buffer. The resin was then washed extensively with lysis buffer. Protein was eluted with Ni-NTA elution buffer (lysis buffer containing an additional 485 mM imidazole). Peak fractions were identified by Bradford assay, pooled, concentrated to ∼2.5 mL with a 3 kDa cutoff spin filter (Millipore), and applied to PD-10 desalting column (Cytiva) pre-equilibrated with lysis buffer to remove imidazole. The elution was incubated with 1 mg TEV protease for 2 hr at 4°C, and injected into a 1 mL HisTrap column (Cytiva) pre-equilibrated with lysis buffer. Peak fractions were identified by Bradford assay, pooled, concentrated to ∼1 mL with the 3 kDa cutoff spin filter (Millipore). Untagged γ-TuNA was then further purified over a HiLoad 16/600 Superdex 75 column pre-equilibrated in gel filtration buffer (40 mM HEPES, pH 7.5, 150 mM NaCl, 1 mM MgCl2 and 2 mM 2-mercaptoethanol). Peak fractions were identified by SDS-PAGE, pooled, concentrated, supplemented with sucrose to 10% (w/v), aliquoted, snap frozen in liquid nitrogen, and stored at –80°C. After TEV cleavage, the resulting CDK5RAP2_44-93_ peptide contains an N-terminal SNIGSGGTPSTGSSLPNVSEE sequence as a remnant of the linking sequence between the TEV cleavage site and the multiple cloning site in the modified expression vector.

To generate the F75A mutant of CDK5RAP2_44-93_, the plasmid above was modified using the Q5 Site-Directed Mutagenesis Kit (NEB) along with the following primers: 5’-GAAAGAAAAC<u>gcg</u>AACCTAAAGCTCCG-3’ and 5’-TTCAATTCAGTGATTTGATTTTC-3’. The plasmid was sequence verified as above and the F75A CDK5RAP2_44-93_ mutant protein was expressed and purified following the same procedure as for wild type CDK5RAP2_44-93_. Both purified CDK5RAP2 fragments were analyzed by SDS-PAGE followed by Coomassie staining (Supplementary Figures S7A and S9B). Protein concentrations were estimated by Bradford assay using BSA as a standard.

### Purification of recombinant MZT2:GST-GCP2-NHD

The coding sequence for MZT2A was PCR amplified from pACEBac1-ZZ-TEV-MZT2-3C-mEGFP (a gift from Tarun Kapoor (Addgene plasmid # 178074 ; http://n2t.net/addgene:178074 ; RRID:Addgene_178074)) using the following primers: 5’-CATCACCCCATGAGCGATTACGACATCCCCACTACTATGGCTATGGCAGGCCTTGC-3’ and 5’-GTGGTGGTGGTGGTGCTCGAGTGCGGCCGCAAGCTTATCAACCGGTGCTGCCCTGCG-3’. The PCR product was subcloned into a pET-M11 vector using restriction-free cloning with Pfu polymerase and subsequent DpnI digestion, resulting in an expression construct for His6-ZZ-TEV-TEV-MZT2.

The coding sequence for amino acids 1-110 of GCP2 were PCR amplified from pACEBac1-gamma-TuSC (a gift from Tarun Kapoor (Addgene plasmid # 178079 ; http://n2t.net/addgene:178079 ; RRID:Addgene_178079)) using the following primers: 5’-GGGGCCCCTGGGATCCATGAGTGAATTTCGGATTCACC-3’ and 5’-GATGCGGCCGCTCGAGTCAGGCTGCAAGCTCAGC-3’. The PCR product was inserted into a BamHI and XhoI double-digested pGEX-6P-1 vector using InFusion Snap Assembly (Takara), resulting in the expression construct for GST-3C-GCP2_1-110_ (referred to as “GST-GCP2-NHD”). Both plasmids were sequence-verified using whole-plasmid sequencing (Microsynth).

To generate GST-tagged MZT2:GCP2-NHD subcomplexes for pull-downs, plasmids for His_6_-ZZ-TEV-MZT2 and GST-GCP2-NHD were co-transformed into BL21-CodonPlus (DE3)-RIL (Stratagene) cells. Proteins were expressed in bacteria grown in 2 L of LB broth after reaching an OD_600_ of 0.6 in the presence of 0.5 mM IPTG at 18 °C and for 16 hr. Cells were harvested by centrifugation at 4000 g at 4 °C for 10 min. Harvested cell pellets were resuspended in ice-cold lysis buffer (20 mM HEPES, pH 7.5, 300 mM NaCl, 15 mM imidazole, and 2 mM 2-mercaptoethanol) and lysed by passing through an Emulsiflex C5 microfuildizer (Avestin) 3 times at >10,000 psi. The lysate was centrifuged at 35,000 rpm in a Type 45 Ti rotor (Beckman) for 1 hr at 4°C. The supernatant was incubated with 1 mL of Ni-NTA agarose (Qiagen) pre-equilibrated with lysis buffer via nutation at 4 °C for 1 hr. The resin was then washed with 50 bed volumes of lysis buffer and protein was eluted with elution buffer (20 mM HEPES, pH 7.5, 150 mM NaCl, 300 mM imidazole, and 2 mM 2-mercaptoethanol). Peak fractions were identified with Bradford assay, pooled and incubated with 1 mg of TEV protease produced in-house, diluted into 1 mL of lysis buffer for 2 hrs and on ice. Digested protein was then diluted 1:1 with GST buffer (20 mM HEPES, pH 7.5, 150 mM NaCl, and 2 mM 2-mercaptoethanol) and incubated with 1 mL Glutathione Sepharose 4 Fast Flow (Cytiva) pre-equilibrated in GST buffer for 1 hr at 4 °C and with nutation. The resin was washed with 20 mL of GST buffer and eluted using GST elution buffer (GST buffer containing 15 mM reduced glutathione). Peak elution fractions were identified using Bradford assay, pooled, concentrated to <250 μL, brought to 10% w/v with sucrose as a cryo-protectant, aliquoted, snap frozen in liquid nitrogen, and stored at -80 °C. The purified protein was analyzed by SDS-PAGE followed by Coomassie staining (Supplementary Figure S7B).

### GST pulldown assays

2 uM of CDK5RAP2_44-93_ with 5 uM of GST or 1 uM, 2 uM, 5 uM, 10 uM of MZT2:GST-GCP-NHD was mixed in binding buffer (40 mM HEPES, 150 mM NaCl, 1 mM MgCl2, 0.1% v/v Tween-20, 1 mM DTT, pH 7.5) and incubated on ice for 1 hr. The mixture was further incubated with glutathione magnetic agarose (Thermo Scientific) pre-equilibrated with binding buffer for 2 hr on ice.The beads were washed twice and resuspended in binding buffer containing SDS-PAGE sample buffer prior to running on gels.

### Purification of recombinant human γ-TuRC (rec-γ-TuRC)

Polycistronic donor plasmid coding for human γ-tubulin, GCP2 and GCP3 (pACEBac1-gamma-TuSC) and human γ-tubulin, GCP2, GCP3, GCP4, GCP5, GCP6, MZT1, ZZ-TEV-MZT2-3C-mEGFP, and actin (pACEBac1-gamma-TuRC-GFP) were a gift from Tarun Kapoor (Addgene plasmid # 178079 ; http://n2t.net/addgene:178079 ; RRID:Addgene_178079). Plasmids were transformed into DH10MultiBacTurbo cells (ATG:biosynthetics GmbH) and transposition-positive colonies were selected and used to generate recombinant bacmids. Bacmids were transfected into Sf9 cells (Novagen) following the Bac-to-Bac manual (Invitrogen), baculoviruses were amplified twice, and fresh virus from γ-TuRC-GFP and γ-TuSC bacmids were mixed together at a 1:1 ratio. This virus mixture was used to infect 1.8–2.4 liters of High Five cells (Thermo Fisher Scientific) at a cell density of 3 × 10^6^ per ml for 60 h at 27°C. Cells were harvested by centrifugation at 1000 g, resuspended in 60 ml ice-cold lysis buffer (40 mM Hepes, pH 7.5, 150 mM KCl, 1 mM MgCl2, 10% glycerol (v/v), 0.1% Tween-20, 0.1 mM ATP, 0.1 mM GTP, 1 mM 2-mercaptoethanol, one cOmplete EDTA-free Protease Inhibitor Cocktail tablets (Roche), and 2 mM PMSF), and lysed by dounce homogenization on ice. The lysate was clarified at 322,000 g for 1 h at 4°C, 0.22-µm syringe–filtered, and loaded onto a 1 mL NHStrap column (Cytiva) previously coupled to 10 mg rabbit IgG (Innovative Biosciences; IRBIGGAP500MG) following the manufacturer’s instructions. The IgG column was washed with lysis buffer followed by gel filtration buffer (40 mM Hepes, pH 7.5, 150 mM KCl, 1 mM MgCl2, 10% glycerol (v/v), 0.1 mM GTP, and 1 mM 2-mercaptoethanol). An expression vector for TEV protease, pRK793, was a gift from David Waugh (Addgene plasmid 8827; http://n2t.net/addgene:8827; Research Resource Identifier: Addgene_8827; Kapust et al., 2001). TEV was expressed in BL21- CodonPlus (DE3)-RIL and purified using Ni-NTA and gel filtration following the methods described in [64]. 1 mg of TEV protease (stored in 40 mM Hepes, pH 7.5, 30% (w/v) glycerol, 150 mM KCl, 1 mM MgCl2, and 3 mM 2-mercaptoethanol) was diluted into 1 ml of gel filtration buffer and injected onto the IgG column, and proteolysis was allowed to proceed for 2 h at 4°C. The digested eluate was pooled, concentrated with a 100 kDa cutoff spin filter (Millipore), and gel filtered over a Superose 6 Increase 10/300 GL column (Cytiva) pre-equilibrated in gel filtration buffer. Peak fractions were pooled and loaded onto two 2-ml sucrose gradient composed of 10%, 20%, 30%, and 40% sucrose (w/v) in gradient buffer (40 mM Hepes, pH 7.5, 150 mM KCl, 1 mM MgCl2, 0.01% Tween-20 (v/v), 0.1 mM GTP, and 1 mM 2-mercaptoethanol). The gradient was centrifuged at 50,000 rpm in a TLS-55 rotor at 4°C for 3 h with minimum acceleration and no break. Fractions were manually collected with a cut-off P1000 pipette tip and analyzed by SDS-PAGE followed by Coomassie staining and/or negative-stain TEM. Peak fractions were aliquoted, snap-frozen, and stored in liquid N2. Gradients were fractionated into 250 µl and analyzed by SDS-PAGE followed by Coomassie staining (Supplementary Figure S9B).

### Biotinylation of anti-GFP nanobody

To biotinylate anti-GFP nanobody, EZ-Link NHS-PEG4-Biotin (Thermo Fisher Scientific) was dissolved in DMSO and added to the concentrated nanobody at a molar ratio of 10:1. The mixture was incubated at 4°C for 4 h and clarified by centrifugation at 21,000 g for 20 min at 4°C. The nanobody was then gel-filtered over a Superdex 75 10/300 column (Cytiva) equilibrated in coupling buffer (100 mM sodium bicarbonate, pH 8.0, and 150 mM NaCl). Biotinylated nanobody was supplemented with 10% glycerol (v/v), flash-frozen in liquid N2, and stored at −80°C.

### Tubulin purification and labeling

Porcine brain tubulin was purified following the method of Castoldi and Popov [65]. Tubulin was covalently labeled with X-rhodamine succinimidyl ester (ThermoFisher Scientific; C6125), Alexa Fluor 488 succinimidyl ester (ThermoFisher Scientific; A20002), or Alexa Fluor 546 succinimidyl ester (ThermoFisher Scientific; A20000) following established methods [66]. Fluorescently-labeled tubulin was typically prepared at a ratio of 0.1-0.2 fluorophores per tubulin dimer. Lyophilized biotinylated porcine tubulin was purchased (Cytoskeleton; T333P) and used at a ratio of 1% for generating biotinylated GMPCPP microtubule seeds.

### Antibodies used in this study

Mouse monoclonal anti-γ-tubulin (clone GTU-88; Merck) at a dilution of 1:1000 was used for the γ-tubulin western blot in Supplementary Figure 1B. HRP-conjugated goat anti-mouse secondary antibody was used at a dilution of 1:10000 for chemiluminescent detection of anti-γ-tubulin antibodies on PVDF membranes.

### Negative staining transmission electron microscopy of S. scrofa γ-TuRC

3.5 µl of sucrose density gradient–purified *S. scrofa* brain γ-TuRC was applied to glow discharged carbon-coated copper grids (EMS; CF400-Cu-50) and incubated for 1 min on ice. Protein solution was removed by manual blotting with Whatman No. 1 filter paper and washed by two drops of 20 ul water and one drop of 20 ul 2% uranyl acetate (w/v). Grids were incubated in another 20 ul uranyl acetate for 45 s. Stain was removed by manual blotting, and grids were air-dried for >24 h in a sealed container layered with desiccant before imaging. Negative stain TEM micrographs of *S. scrofa* brain γ-TuRC were collected on a TFS Morgagni 268 microscope operating at 100 kV. The representative micrograph was collected at 20,000X.

### Cryo-electron microscopy of the *S. scrofa* γ-TuRC

EM grids were prepared by overlaying Quantifoil 300 mesh (copper, R 2/2) holy carbon grids with homemade continuous carbon film. The grids were glow discharged at 15 mA for 15 s and placed on an ice cold block via metal forceps. 2.5 μL of thawed γ-TuRC sample was applied to the grid for 5 min and manually blotted away. Another 2.5 μL of sample was replaced on the grid and this procedure was repeated for up to 8 applications. After the final application, the grid was washed by incubating twice with 20 μL of washing buffer (40 mM HEPES, pH 7.5, 150 mM NaCl, 1 mM MgCl2, 1 mM EGTA, 2 mM 2-mercaptoethanol, 0.01% IGEPAL, 0.1 mM GTP) for 1 min. The grid was supplemented with an additional 3.5 μL of applied washing buffer, transferred to a Vitrobot (ThermoFisher Scientific), and blotted for 4 s at 100% humidity and 4°C, then plunge-frozen into liquid ethane and stored in liquid nitrogen.

109,725 movies were collected with a Gatan K3, CDS mode, and a slit width of 15-20 eV on a GIF BioQuantum energy filter in a TFS Titan Krios G3i (2) FEG of the ScopeM facility at ETH (Zürich, Switzerland). Automatic data collection was performed with “Faster acquisition mode” in EPU software (Thermo Fisher Scientific). Full data collection statistics are listed in Supplementary Table S2.

### Image processing

For all data sets a random selection of 6,000 movies per group were used to estimate a gain reference of each acquisition with relion_estimate_gain [67]. .mrc movie files were imported in cryoSPARC v4.3.x - v4.4.0 for Patch Motion Correction and Patch CTF estimation [45]. An initial attempt using templates for picking was initiated with EMD-21073 [10]. Subsequently, the extracted particles underwent 2D classification. During this analysis, a distinctive ring-shaped class corresponding to the γ-TuRC emerged. An extra picking step was initiated with TOPAZ [68], a TOPAZ Cross Validation in cryoSPARC allowed us to explore the parameters for Training a model over 495 micrographs from Datasets 1, 2 and 3. We generated models for independent datasets and TOPAZ extract functions. 2D classifications were performed and good particles were selected for 3D reconstructions. 3D refinements and reconstructions were done in cryoSPARC, RELION 4 and cryoDRGN 1.1.0 [10,45]. Particles files were moved from cryoSPARC to RELION with UCSF PyEM [69].

Independently, 3 main classes of γ-TuRCs: Open, Partially-assembled and CMG-decorated class were obtained in cryoSPARC and RELION. A subset of good particles were simultaneously 3D classified (force hard classification) and refined in 384 box size volumes with the 3 initial γ-TuRC like structures and a “trash collector”. Multiple runs of focus 3D classification and focus/local 3D reconstructions were performed in cryoSPARC and RELION. To further explore the heterogeneity of the partially-assembled particles containing cryoSPARC, poses were imported in cryoDRGN. Soft masks were inspected with UCSF Chimera or ChimeraX [70,71]. External function Sidesplitter was applied during 3D classification and/or 3D auto-refine in RELION 4 [72]. A selection of maps from the cryo-EM image processing pipeline were used for model building the different γ-TuRC subclasses of this research (Supplementary Figure S6 and Supplementary Table S3).

### Model building

The *S. scrofa* γ-tubulin (A0A287BRH5) model was built using homology modeling through the SwissModel server [73]. For GCP2 (A0A480VJI0), GCP3 (F1RN46), MZT2 (F1RK97), and CDK5RAP2, as well as N-terminal (amino acids 1 - 124) GCP5 (I3L738) and MZT1 (A0A287AS02), we employed AlphaFold2 to generate initial models [40]. Models for GCP4 (F1SI61), GCP5 (I3L738), GCP6 (F1RXS1), and MZT1 (A0A287AS02) were obtained with either Phyre2 and AlphaFold2 [74].

Subsequently, we visualized and aligned the individual components in UCSF Chimera and ChimeraX [70,71]. The “fit in map” function was used to rigid-body fit the components into their respective positions within the consensus density maps for each γ-TuRC conformation: open, partially-assembled, and CMG-decorated. All model components were further rigid-body fit and visually inspected into their corresponding focus maps generated using the “Chain-by-chain” rigid body fitting feature in Coot 0.9.8 [75].

Then, rigid body-fitted components in their local maps underwent molecular dynamics flexible fitting (MDFF) in VMD v1.9.3 [76]. The MDFF run was included during a *Namdinator* job with NAMD v2.14, which comprised 2,000 minimization steps, 20,000 simulation steps in a vacuum, and applied a grid force of 0.3 [77]. Flexibly fitted models were then subjected to a Phenix v1.20.1-4487 real-space refinement including 5 macrocycles [78].

Upon further visual examination utilizing Coot and ChimeraX, we identified and selected the optimal chain and regions for individual components refined in local maps. This process involved trimming disordered regions in models lacking corresponding density in individual maps. Lastly, models went through a second *Namdinator* step that included 5 macrocycles of a Phenix real-space refinement using maps for the open, partially-assembled and CMG-decorated classes which were low pass filtered to 8 Å using “relion_image_handler”. Model refinement statistics are provided in Supplementary Table S4.

Free energy calculations between specific γ-TuRC subunit interfaces were calculated using the PRODIGY web server [50].

### Mass spectrometry

Proteins in 20 µL solution were precipitated with trichloroacetic acid (TCA; Sigma-Aldrich) at a final concentration of 5% and washed twice with ice-cold acetone. Protein pellets were air dried and re-solubilized in 45 µl of digestion buffer (10 mM Tris, 2 mM CaCl2, pH 8.2). 500 ng of Sequencing Grade Trypsin (Promega) were added for digestion carried out in a microwave instrument (Discover System, CEM) for 30 min at 5 W and 60 °C. The samples were dried to completeness and re-solubilized in 20 µL of MS sample buffer (3% acetonitrile, 0.1% formic acid). Peptide concentration was determined using the Lunatic UV/Vis polychromatic spectrophotometer (Unchained Labs).

Mass spectrometry analysis was performed on an Orbitrap Exploris 480 mass spectrometer (Thermo Fisher Scientific) equipped with a Nanospray Flex Ion Source (Thermo Fisher Scientific) and coupled to an M-Class UPLC (Waters). Solvent composition at the two channels was 0.1% formic acid for channel A and 0.1% formic acid, 99.9% acetonitrile for channel B. Column temperature was 50°C. For each sample 8 µL of peptides were loaded on a commercial nanoEase MZ Symmetry C18 Trap Column (100 Å, 5 µm, 180 µm x 20 mm, Waters) followed by a nanoEase MZ C18 HSS T3 Column (100 Å, 1.8 µm, 75 µm x 250 mm, Waters). The peptides were eluted at a flow rate of 300 nL/min. After a 3 min initial hold at 5% B, a gradient from 5 to 22 % B in 40 min and 22 to 32% B in additional 5 min was applied. The column was cleaned after the run by increasing to 95 % B and holding 95 % B for 10 min prior to re-establishing loading condition for another 10 minutes.

LC-MS/MS analysis was performed on an Q Exactive mass spectrometer (Thermo Scientific) equipped with a Digital PicoView source (New Objective) and coupled to an M-Class UPLC (Waters). Solvent composition at the two channels was 0.1% formic acid for channel A and 0.1% formic acid, 99.9% acetonitrile for channel B. For each sample, 2 µl of peptides were loaded on a commercial ACQUITY UPLC M-Class Symmetry C18 Trap Column (100 Å, 5 µm, 180 µm x 20 mm, Waters) connected to a ACQUITY UPLC M-Class HSS T3 Column (100 Å, 1.8 µm, 75 µm X 250 mm, Waters). The peptides were eluted at a flow rate of 300 nL/min. After a 3 min initial hold at 5% B, a gradient from 5 to 35 % B in 42 min and 35 to 40% B in additional 5 min was applied. The column was cleaned after the run by increasing to 95 % B and holding 95 % B for 10 min prior to re-establishing loading condition.

The mass spectrometer was operated in data-dependent mode (DDA) using Xcalibur, with spray voltage set to 2.3 kV and heated capillary temperature at 275 °C. Full-scan MS spectra (350−1500 m/z) were acquired at a resolution of 70’000 at 200 m/z after accumulation to a target value of 3,000,000, followed by HCD (higher-energy collision dissociation) fragmentation on the twelve most intense signals per cycle. Ions were isolated with a 1.2 m/z isolation window and fragmented by higher-energy collisional dissociation (HCD) using a normalized collision energy of 25 %. HCD spectra were acquired at a resolution of 35,000 and a maximum injection time of 120 ms. The automatic gain control (AGC) was set to 100,000 ions. Charge state screening was enabled and singly and unassigned charge states were rejected. Only precursors with intensity above 25,000 were selected for MS/MS. Precursor masses previously selected for MS/MS measurement were excluded from further selection for 20 s, and the exclusion window tolerance was set at 10 ppm. The samples were acquired using internal lock mass calibration on m/z 371.1010 and 445.1200.

The mass spectrometry proteomics data were handled using the local laboratory information management system (LIMS) [70]. The acquired raw MS data were processed by MaxQuant (version 2.0.1), followed by protein identification using the integrated Andromeda search engine [71]. Spectra were searched against the Uniprot Pig reference proteome (taxonomy 9823, canonical version from 2019-12-03) or Uniprot human reference proteome (taxonomy 9606, canonical version from 2023-03-30), concatenated to its reversed decoyed fasta database and common protein contaminants. Methionine oxidation, Deamidation (NQ) and N-terminal protein acetylation were set as variable modifications and Carbamidomethylation (C) was set as fixed modification. Enzyme specificity was set to trypsin/P allowing a minimal peptide length of 7 amino acids and a maximum of two missed-cleavages. MaxQuant Orbitrap default search settings were used. The maximum false discovery rate (FDR) was set to 0.01 for peptides and 0.05 for proteins. Label free quantification was enabled and a 2 minute window for match between runs was applied. In the MaxQuant experimental design template, each file is kept separate in the experimental design to obtain individual quantitative values. Scaffold (Proteome Software Inc., version 5.10) was used to validate MS/MS based peptide and protein identifications. Peptide identifications were accepted if they achieved a false discovery rate (FDR) of less than 0.1% by the Scaffold Local FDR algorithm. Protein identifications were accepted if they achieved an FDR of less than 1.0 % and contained at least 2 identified peptides.

### Total internal reflection fluorescence microscopy

The microscope setup was built around a Nikon N-STORM microscope equipped with an SR HP Apo TIRF 100xH objective with a numerical aperture of 1.49, operated under immersion oil (Nikon), three-axis piezo-electric stages MCL NanoDrive PiezoZ Drive, an Andor DU-897 EM-CCD or a Hamamatsu Orca Flash 4 v3 sCMOS camera, and two lasers generating 488 nm and 561 nm excitation light. Image acquisition was controlled using NIS Elements AR 5.21.01 (Nikon).

Glass slides (Premiere 8201) and glass coverslips (Corning, 22 × 22 mm, 2845-22) were washed for 5 min in acetone, rinsed with water, sonicated for 10 min in 50 % methanol, rinsed with water, sonicated for 10 minutes in 0.5 M KOH and rinsed with water before drying using filtered compressed air. Then, the clean, dry glass was subjected to plasma cleaning for 3 minutes at high intensity in a Harrick Plasma PDC-32G-2 plasma cleaner connected to an ICME M71B4 vacuum pump. Coverslips were functionalized by silanization with 0.1% dichlorodimethylsilane (v/v) in heptane for 2 hours. For experiments in Figure 5 and Supplementary Figure S9, glass slides were also silanized. After silanization, glass was submerged in clean heptane, sonicated for 20 minutes, and then washed in water and dried using filtered compressed air. Silanized coverslips were attached to glass slides using two strips of double-sided tape (3M) separated by ∼5 mm.

For experiments in Supplementary Figure S1, the channel was rinsed with BRB80 buffer, followed by the application of *S. scrofa* brain γ-TuRC diluted in BRB80 or an empty BRB80 buffer as a control. After 5 minutes of incubation, non-adherent molecules were washed away by BRB80, and the channel was treated for 5 minutes with 1% Pluronic F127 (Sigma Aldrich) in BRB80. The channel was washed with an assay buffer containing BRB80 + 1 mM DTT (Axon Lab AG), 50 mM KCl, 0.15% (w/v) methylcellulose (Sigma Aldrich), 0.2 mg/ml BSA (Axon Lab AG), and 1 mM GTP (Jena Bioscience). A 12.5 µM solution of fluorescently labeled tubulin in 1x assay buffer containing an oxygen scavenger system (freshly mixed 0.035 mg/ml catalase (Sigma Aldrich), 0.2 mg/ml glucose oxidase (Merck), 2.5 mM glucose (Sigma Aldrich), and 10 mM DTT) was introduced to the flow cell. Immediately afterwards, the flow cell was sealed with nail polish and placed on the stage of the TIRF microscope, which was equilibrated to 37 °C prior to measurement. Data were collected for an overall period of 10 minutes in ∼16-second intervals for the red channel (excitation at 561 nm) with 100 ms exposure. For each experimental condition, 10 fields of view per biological repetition and three independent repetitions were collected. Image stacks were drift-corrected in FIJI [79] using the MultiStackReg plugin created by Brad Busse (https://github.com/miura/MultiStackRegistration).

For experiments in Supplementary Figure S9, the channel was rinsed with BRB80 buffer followed by the application of *S. scrofa* brain γ-TuRC diluted in BRB80 or an empty BRB80 buffer as a control. After 5 minutes of incubation, non-adherent molecules were washed away by BRB80, and the channel was treated for 5 minutes with 1% Pluronic F127 in BRB80. The channel was washed with the same assay buffer as introduced before. A 12.5 µM solution of fluorescently labeled tubulin in 1x assay buffer containing an oxygen scavenger system as explained above and when applicable CDK5RAP2_44-93_ was introduced to the flow cell. Immediately afterwards, the flow cell was sealed with VALAP (1:1:1 vaseline, linoleum, paraffin wax) and placed on the preheated (35°C) microscope stage. Each sample was measured for 22 minutes in 10 s intervals with excitation at 561 nm (100 ms exposure) and 488 nm (400 ms exposure). Three fields of view were collected in every channel. Microtubules were counted manually at 21 minutes after insertion of the sample in FIJI [79].

For experiments with rec-γ-TuRC in Figure 5, the channel was rinsed with BRB80 buffer, followed by biotinylated BSA (Life Technologies Europe) diluted in BRB80. After 5 minutes of incubation, non-adherent molecules were washed away by BRB80, and the channel was treated for 5 minutes with 1% Pluronic F127 in BRB80. The channel was washed with BRB80 and then incubated with a neutravidin solution containing 0.25 mg/mL Neutravidin (Thermo Scientific), 1 mg/mL BSA, 0.04%Tween20 (Sigma Aldrich) and 10 mM DTT. The channel was washed with BRB80 followed by 5 minutes incubation with biotinylated GFP nanobody (described above). The channel was washed with the assay buffer mentioned above, followed by incubating the channel with a 1:10 (v/v) rec-γ-TuRC solution for 5 minutes. After washing the channel again with assay buffer, a 12.5 µM solution of fluorescently labeled tubulin in assay buffer containing an oxygen scavenger system as mentioned above and when applicable additional CDK5RAP2_44-93_ was introduced to the flow cell. Immediately afterwards, the flow cell was sealed with VALAP and placed on the preheated (35°C) microscope stage. Measurement settings and analysis methods were the same as explained above for Supplementary Figure S9. GFP puncta were counted using an in-house written python script which includes background removal and a spot detection package from scikit-image.

It should be noted that the S. scrofa and rec-γ-TuRC CDK5RAP244-93 activation experiments are difficult to directly compare, as they employ distinct surface-immobilization strategies, the concentration of the native material is typically much lower than for rec-γ-TuRC, and the S. scrofa γ-TuRC here is not fluorescent, so the final number adhered on the coverslip surface was not determined.

### Helical parameter analysis

An oligomeric structure of the *S. cerevisiae* γ-TuRC in the closed conformation was manually built from an available γ-TuSC subunit (PDB ID: 5FLZ [29]). Seven neighbouring copies of the γ-TuSC was docked into a density map for an engineered γ-TuSC oligomer in the closed conformation (EMD-2799 [20]) using PyMoL (The PyMOL Molecular Graphics System, Version 2.0 Schrödinger, LLC.) and Chimera softwares. γ-TuRC structures, including those generated in this study, were then aligned to the *S. cerevisiae* closed γ-TuRC model via their GRIP1 domains using the align command in PyMoL. The aligned structures were subsequently read out into PDB format and analyzed using a custom python script. The aligned PDB files were loaded and coordinates of all Cα atoms in the structures were extracted using biophython [80]. The position of each γ-tubulin was given by the center of mass (CoM), i.e., the mean of its Cα atoms. The CoM was used to analyze structural changes due to the robustness and stability concerning conformational changes of individual γ-tubulins. The Euclidean distance between the CoM of the *S. cerevisiae* γ-TuRC in the closed conformation and all other γ-TuRC structures at each position was calculated to analyze changes between complexes.

To examine changes in orientation of individual γ-tubulins across the structures, the CoM of the Cαs of the first half and second half of γ-tubulin helix 12 was used to define a vector describing tilt and rotation of each γ-tubulin. The coordinates were transformed to spherical coordinates, where phi (φ) denotes the angle of the vectors projected in x-y-plane, while theta (Ө) describes the tilt away from the z-axis (the length of the vectors are set to 1 for simplicity). Normalization of theta and phi was done by subtraction, resulting in the direction of the change in orientation of each γ-tubulin with respect to the *S. cerevisiae* closed, pseudo-γ-TuRC reference structure.

## Code availability

Custom scripts for TIRF image GFP punctum detection and microtubule background subtraction, as well as the helical analysis of γ-TuRC structures, will be made available via GitHub.

