## Supplementary Information for "Closure of the γ-tubulin ring complex by CDK5RAP2 activates microtubule nucleation"

**Supplementary Figures**

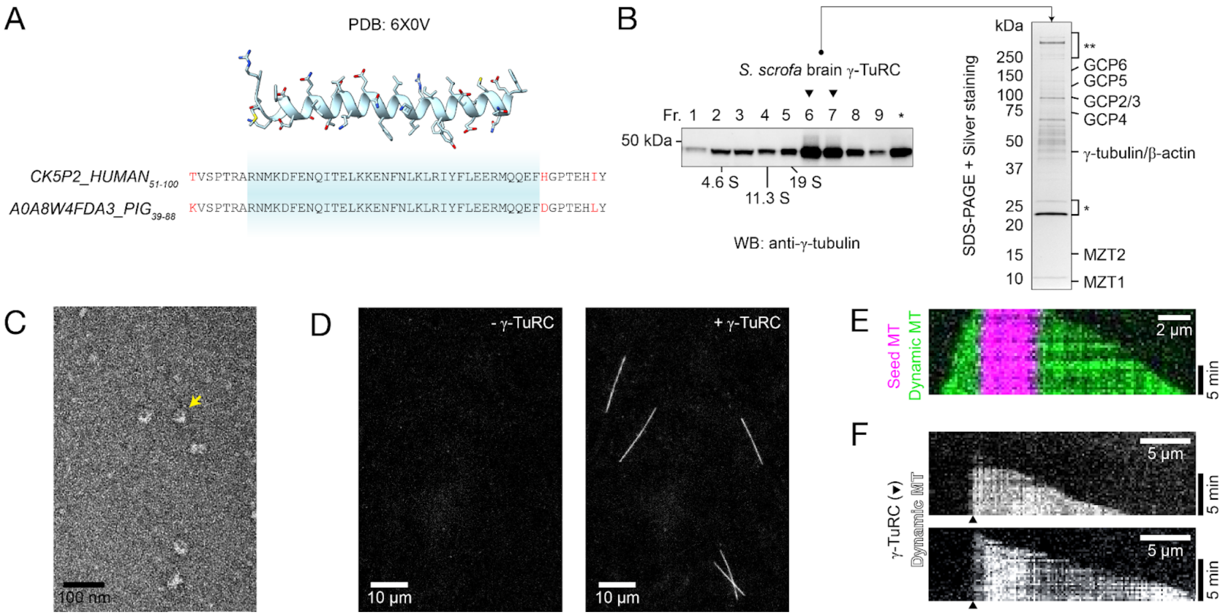


**Figure S1. Isolation and characterization of *S. scrofa* brain γ-TuRC.** A) Sequence alignment between CDK5RAP2$_{51\text{-}100}$ (referred to as CK5P2_HUMAN in the Uniprot database) and the homologous sequence from *S. scrofa* (Accession number: A0A8W4FDA3_PIG). Non-identical residues are coloured in red. The sequence for the previously-resolved model for the CDK5RAP2 is highlighted in cyan, and a cartoon and stick representation of the molecular model is shown above for reference [1]. B) Left: Anti-γ-tubulin Western blot of *S. scrofa* brain γ-TuRC sucrose density gradient fractions. Peak γ-TuRC fractions used for subsequent studies are indicated by black triangles. Standards with known S-values (BSA (4.6 S), catalase (11.3 S) and thyroglobulin (19 S)) were run in parallel (not shown); the corresponding peak fractions are also indicated. The last fraction corresponds to a presumed mixture of pellet/precipitated complexes and is indicated by an asterisk. Right: Silver-staining SDS-PAGE analysis of *S. scrofa* brain γ-TuRC. γ-TuRC components identified by LC-MS/MS are indicated according to their predicted molecular weight. Contaminations likely corresponding to residual 3C protease and/or uncleaved GFP-CDK5RAP2$_{51\text{-}100}$ (*) as well as dynein and/or β-spectrin (**) are indicated. C) Transmission EM micrograph of negatively stained *S. scrofa* γ-TuRC. Yellow arrow indicates an example γ-TuRC particle. D) Total internal reflection fluorescence (TIRF) microscopy images of microtubule nucleation reactions with surfaces treated without (left) and with (right) γ-TuRC. On average, the *S. scrofa* γ-TuRC nucleation rate was low, with 1 microtubule observed in every ~7 fields of view collected; however, no microtubules were detected in the absence of surface-adhered *S. scrofa* γ-TuRC (n = 46 fields of view from 5 independent experiments performed on 2 separate days). E) Two-color kymograph from a time-lapse sequence of a dynamic microtubule (“Dynamic MT”; green) nucleated by a stabilized seed (“seed MT”; magenta). F) Two example kymographs of a dynamic microtubule nucleated by *S. scrofa* γ-TuRC (not fluorescent, but its presumed position is marked by a black triangle). The average growth rate of microtubules nucleated by surface-adhered *S. scrofa* γ-TuRC was 2.4 ± 0.25 μm min$^{-1}$ (n = 10 microtubules from 3 experiments).

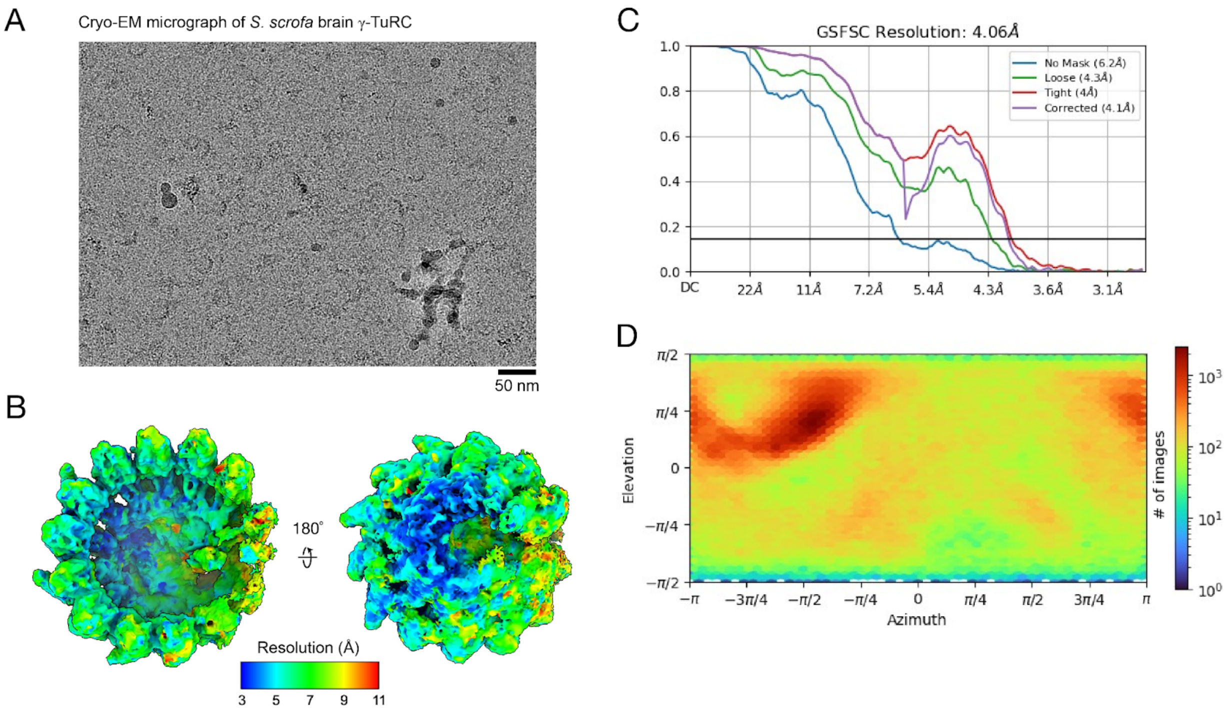


**Figure S2. Consensus cryo-EM reconstruction of *S. scrofa* brain γ-TuRC.** A) Cryo-EM micrograph of *S. scrofa* brain γ-TuRC. B) Two views of the overall *S. scrofa* γ-TuRC density map analyzed by ResMap [2], showing a resolution distribution ranging from 3 to >11 Å. C) Gold-standard Fourier shell correlation plot for the overall *S. scrofa* brain γ-TuRC density map. D) Euler angle orientation distribution for *S. scrofa* brain γ-TuRC particles. Panels C) and D) were generated in cryoSPARC [3].

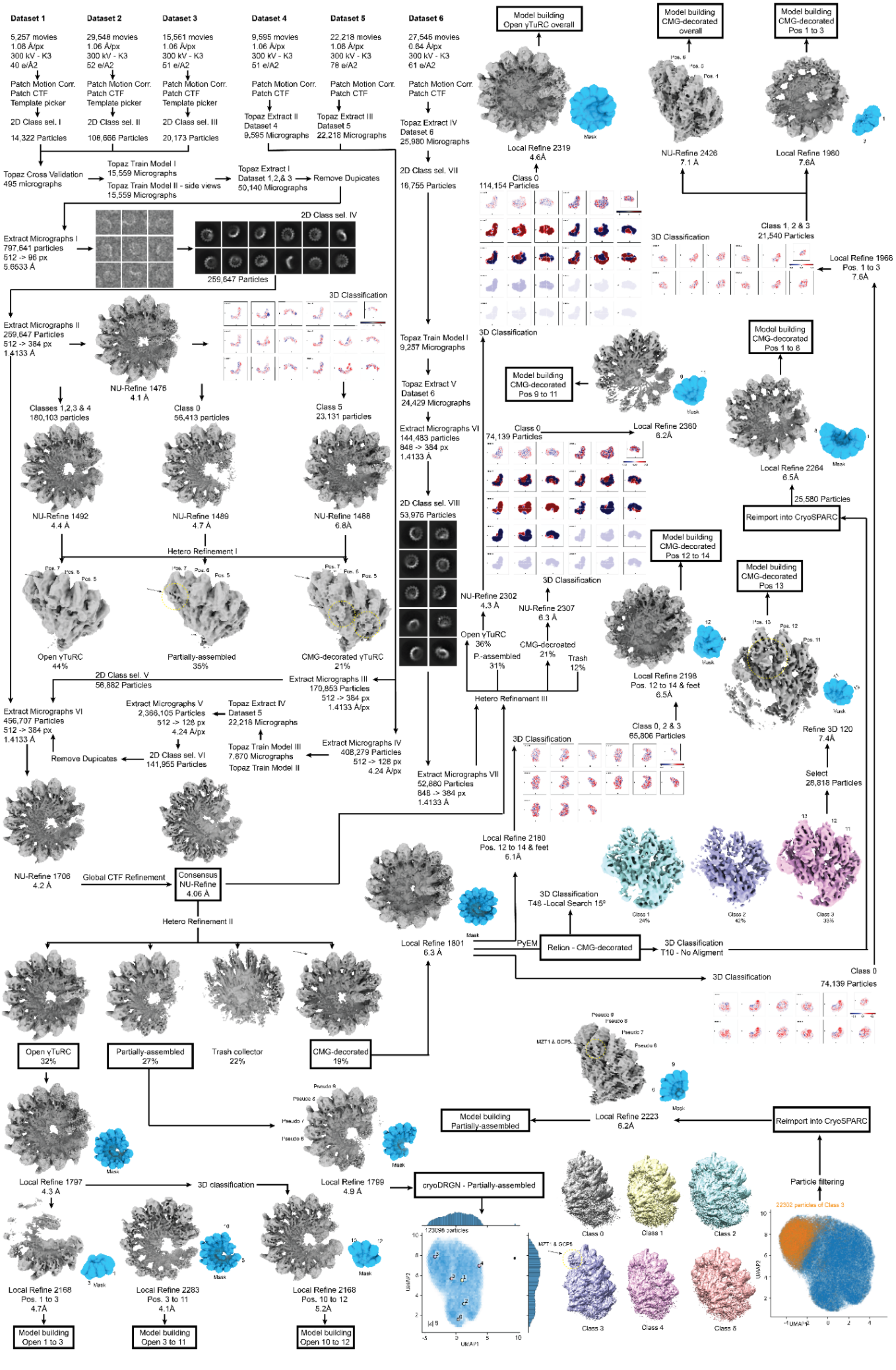
Dataset 1
5,257 movies
1.06 Å/px
300 kV - K3
40 e/Å2
Patch Motion Corr.
Patch CTF
Template picker
2D Class sel. I
14,322 Particles
Dataset 2
29,548 movies
1.06 Å/px
300 kV - K3
52 e/A2
Patch Motion Corr.
Patch CTF
Template picker
2D Class sel. II
108,666 Particles
Dataset 3
15,561 movies
1.06 Å/px
300 kV - K3
51 e/A2
Patch Motion Corr.
Patch CTF
Template picker
2D Class sel. III
20,173 Particles
Dataset 4
9,595 movies
1.06 Å/px
300 kV - K3
51 e/A2
Patch Motion Corr.
Patch CTF
Topaz Extract II
Dataset 4
9,595 Micrographs
Dataset 5
22,218 movies
1.06 Å/px
300 kV - K3
78 e/A2
Patch Motion Corr.
Patch CTF
Topaz Extract III
Dataset 5
22,218 Micrographs
Dataset 6
27,546 movies
0.64 Å/px
300 kV - K3
61 e/A2
Patch Motion Corr.
Patch CTF
Topaz Extract IV
Dataset 6
25,980 Micrographs
2D Class sel. VII
16,755 Particles
Topaz Cross Validation
495 micrographs
Topaz Train Model I
15,559 Micrographs
Topaz Train Model II - side views
15,559 Micrographs
Topaz Extract I
Dataset 1,2,& 3
50,140 Micrographs
Remove Dupicates
Extract Micrographs I
797,641 particles
512 -> 96 px
5.6533 Å
2D Class sel. IV
259,647 Particles
Extract Micrographs II
259,647 Particles
512 -> 384 px
1.4133 Å
NU-Refine 1476
4.1 Å
3D Classification
Classes 1,2,3 & 4
180,103 particles
Class 0
56,413 particles
Class 5
23,131 particles
NU-Refine 1492
4.4 Å
NU-Refine 1489
4.7 Å
NU-Refine 1488
6.8Å
Hetero Refinement I
Open γTuRC
44%
Partially-assembled
35%
CMG-decorated γTuRC
21%
2D Class sel. V
56,882 Particles
Extract Micrographs III
170,853 Particles
512 -> 384 px
1.4133 Å/px
Extract Micrographs VI
456,707 Particles
512 -> 384 px
1.4133 Å
Extract Micrographs V
2,366,105 Particles
512 -> 128 px
4.24 Å/px
Topaz Extract IV
Dataset 5
22,218 Micrographs
Remove Dupicates
2D Class sel. VI
141,955 Particles
Topaz Train Model III
7,870 Micrographs
Topaz Train Model II
Extract Micrographs IV
408,279 Particles
512 -> 128 px
4.24 Å/px
NU-Refine 1706
4.2 Å
Global CTF Refinement
Consensus
NU-Refine
4.06 Å
Hetero Refinement II
Open γTuRC
32%
Partially-assembled
27%
Trash collector
22%
CMG-decorated
19%
Local Refine 1797
4.3 Å
3D classification
Local Refine 1799
4.9 Å
cryoDRGN - Partially-assembled
Local Refine 2168
Pos. 1 to 3
4.7Å
Local Refine 2283
Pos. 3 to 11
4.1Å
Local Refine 2168
Pos. 10 to 12
5.2Å
Model building
Open 1 to 3
Model building
Open 3 to 11
Model building
Open 10 to 12
Topaz Train Model I
9,257 Micrographs
Topaz Extract V
Dataset 6
24,429 Micrographs
Extract Micrographs VI
144,483 particles
848 -> 384 px
1.4133 Å
2D Class sel. VIII
53,976 Particles
Extract Micrographs VII
52,880 Particles
848 -> 384 px
1.4133 Å
Model building
Open γTuRC overall
Local Refine 2319
4.6Å
Class 0
114,154 Particles
3D Classification
Model building
CMG-decorated
Pos 9 to 11
Class 0
74,139 Particles
Local Refine 2360
6.2Å
3D Classification
NU-Refine 2302
4.3 Å
NU-Refine 2307
6.3 Å
Open γTuRC
36%
P.-assembled
31%
CMG-decroated
21%
Trash
12%
Hetero Refinement III
Model building
CMG-decorated
overall
NU-Refine 2426
7.1 Å
Model building
CMG-decorated
Pos 1 to 3
Local Refine 1980
7.6Å
Class 1, 2 & 3
21,540 Particles
3D Classification
Local Refine 1966
Pos. 1 to 3
7.6Å
Model building
CMG-decorated
Pos 1 to 6
Local Refine 2264
6.5Å
25,580 Particles
Reimport into CryoSPARC
Model building
CMG-decorated
Pos 12 to 14
Local Refine 2198
Pos. 12 to 14 & feet
6.5Å
Model building
CMG-decorated
Pos 13
Refine 3D 120
7.4Å
Select
28,818 Particles
Class 0, 2 & 3
65,806 Particles
3D Classification
Local Refine 2180
Pos. 12 to 14 & feet
6.1Å
3D Classification
T48 -Local Search 15º
Class 1
24%
Class 2
42%
Class 3
33%
Local Refine 1801
6.3 Å
PyEM
Relion - CMG-decorated
3D Classification
T10 - No Aligment
3D Classification
Class 0
74,139 Particles
Model building
Partially-assembled
Local Refine 2223
6.2Å
Reimport into CryoSPARC
Particle filtering
22302 particles of Class 3
Class 0
Class 1
Class 2
Class 3
Class 4
Class 5
UMAP1
UMAP2
Mask

**Figure S3. Data processing workflow for reconstruction of γ-TuRC density maps**.

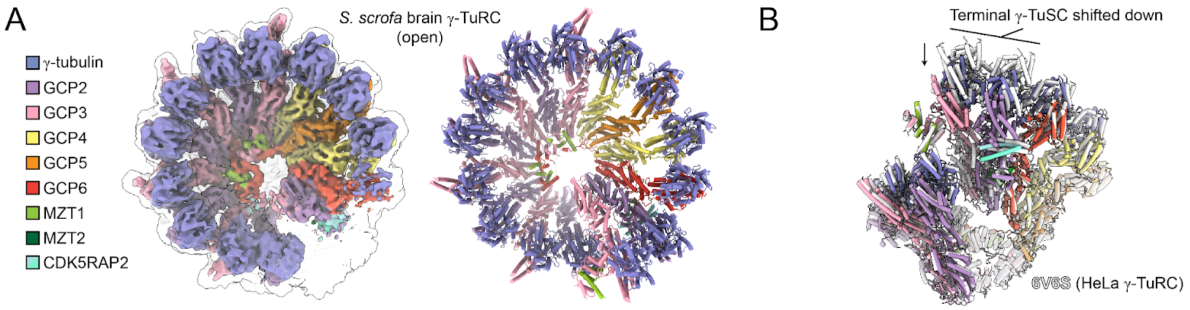


**Figure S4. Structure of the *S. scrofa* γ-TuRC in the open conformation.** A) Left: Surface representation top view of a sharpened density map for the 3D classified *S. scrofa* γ-TuRC in the open conformation. An 8 Å low-pass filtered, unsharpened map is shown in transparent white surface to show density for the terminal γ-TuSC. Right: Cartoon representation top view of a model for the *S. scrofa* γ-TuRC in the open conformation, coloured according to the legend on the left. B) Tilted side view focusing on the γ-TuRC "seam" region of the human CM1-purified γ-TuRC (white cartoon representation; PDB ID: 6V6S [4]), aligned to the *S. scrofa* γ-TuRC in the open conformation (cartoon representation coloured according to the legend in A)). The downward-shifted terminal γ-TuSC is indicated.

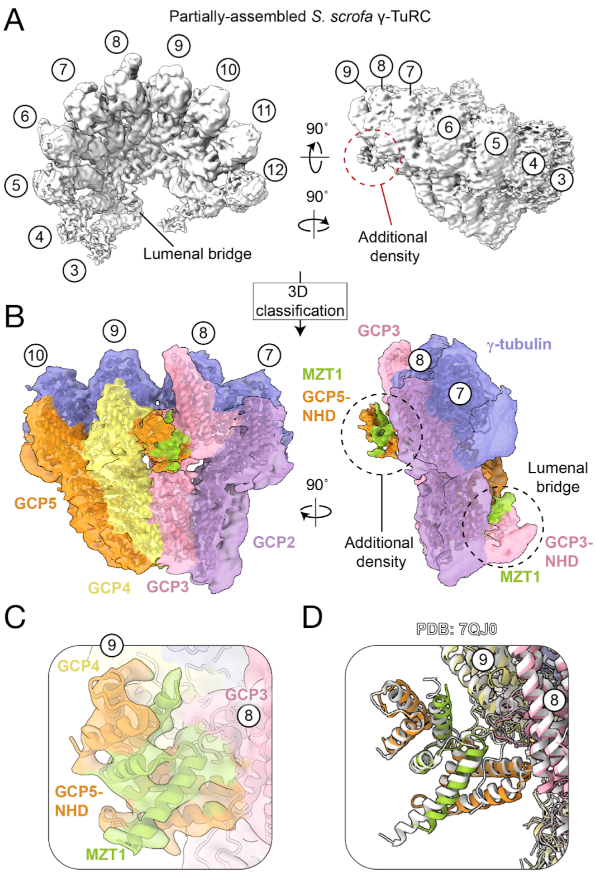


**Figure S5. Structure of the *S. scrofa* MZT1/GCP5-NHD module in the partially-assembled γ-TuRC subclass.** A) Two views of the *S. scrofa* brain γ-TuRC "partially-assembled" density map. Pseudo-positions 3-12 of the γ-TuRC, as well as the lumenal bridge, are indicated. An additional density situated between positions 8 and 9 on the outer face of the complex observed at lower contour levels is highlighted by a red dashed circle. B) Two views of a 3D classified density map (transparent coloured surface representation), showing pseudo-positions 7-10 of the partially-assembled γ-TuRC. The left image shows the view from the outer face of the complex. The additional density is better-resolved and is highlighted by a dashed circle, as is the MZT1:GCP3-NHD module in the lumenal bridge. Corresponding refined models for γ-tubulin, GCP2, GCP3, GCP4, GCP5, and MZT1 are shown in cartoon representation and are coloured according to the legend in Figure 1. C) Close up view of the MZT1/GCP5-NHD density (transparent coloured surface representation) with the corresponding γ-TuRC models shown in cartoon representation. Positions 8 and 9 are indicated. D) Cartoon representations of aligned models from this study (coloured) and the previously reported MZT1:GCP5-NHD-decorated 6-spoke assembly intermediate (PDB ID: 7QJ0 [5]; RMSD for MZT1:GCP5-NHD = 1.1 Å).

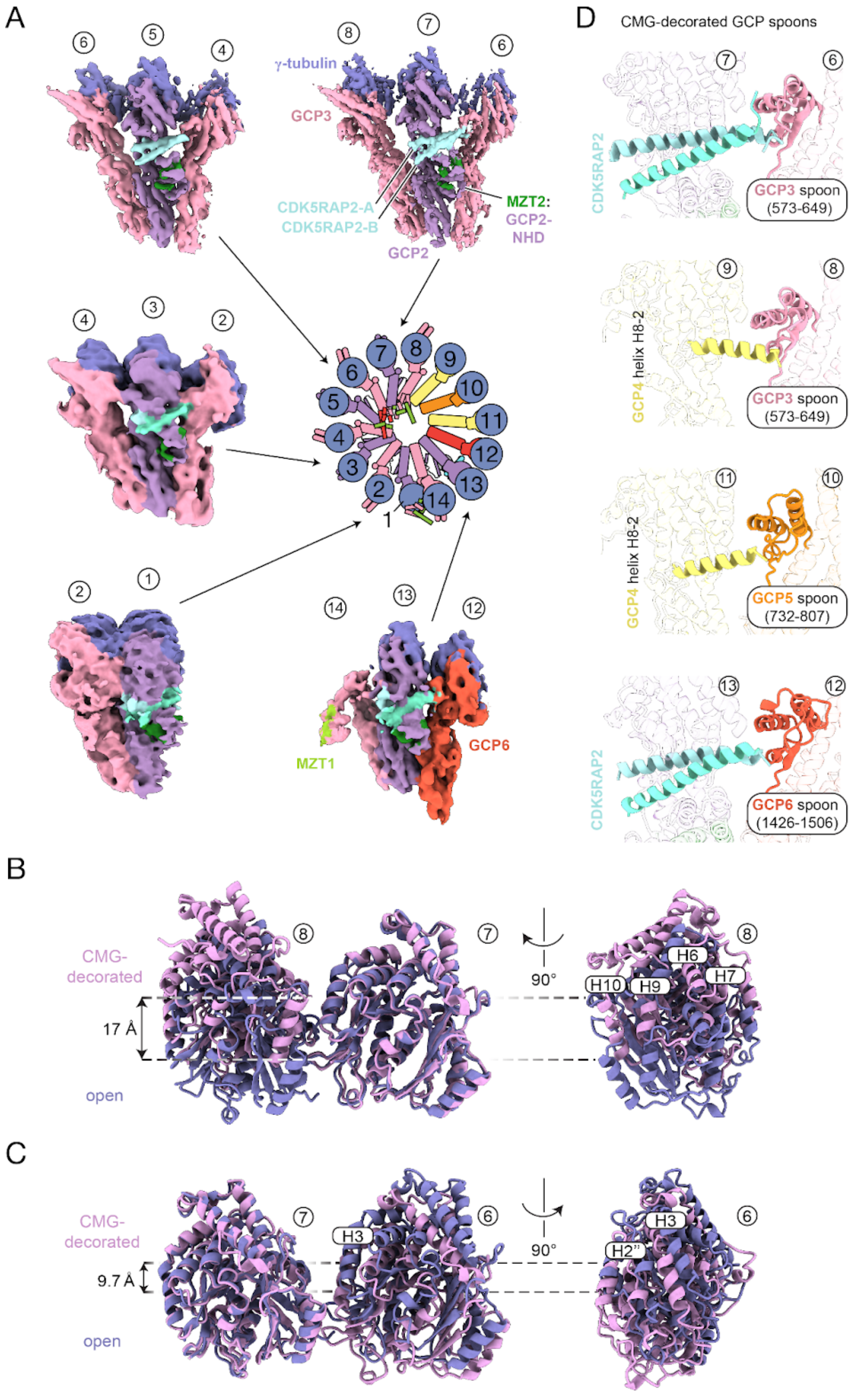
A
γ-tubulin
GCP3
CDK5RAP2-A
CDK5RAP2-B
MZT2:
GCP2-
NHD
GCP2
MZT1
GCP6
D
CMG-decorated GCP spoons
CDK5RAP2
GCP3 spoon (573-649)
GCP4 helix H8-2
GCP3 spoon (573-649)
GCP4 helix H8-2
GCP5 spoon (732-807)
CDK5RAP2
GCP6 spoon (1426-1506)
B
CMG-decorated
17 Å
open
90°
H6
H7
H10
H9
C
CMG-decorated
9.7 Å
open
H3
90°
H3
H2''

**Figure S6. Overview of CMG densities, sequence alignment of MZT2:GCP2-NHD from *H. sapiens* and *S. scrofa*, γ-tubulin:γ-tubulin interface details, and GCP spoon binding interfaces.** A) Surface representations of zoned density maps for each of the CMG modules in the CMG-decorated γ-TuRC, coloured according to the legend in Figure 2. Views are all from the outer face of the complex, and γ-TuRC positions are indicated above each γ-tubulin. A schematic of the γ-TuRC is shown in the center for reference. B) Two views of γ-tubulins from positions 8 and 7 of the open (violet) and CMG-decorated (closed, pink) conformation, aligned at position 7 (RMSD = 1.2 Å). α-helices are labeled and a ~17 Å displacement is indicated. In the right view, the aligned γ-tubulins are omitted for clarity. C) Two views of γ-tubulins from positions 7 and 6 of the open (violet) and CMG-decorated (closed, pink) conformation, aligned at position 7 (RMSD = 1.2 Å). α-helices are labeled and a ~9.7 Å displacement is indicated. In the right view, the aligned γ-tubulins are omitted for clarity. A-helices mentioned in the text are indicated. D) Cartoon representations of elements in the CMG-decorated γ-TuRC contacting GCP spoons at positions 6, 8, 10, and 12. The rest of the GCP structures are also shown in cartoon representations but are dimmed in colour.

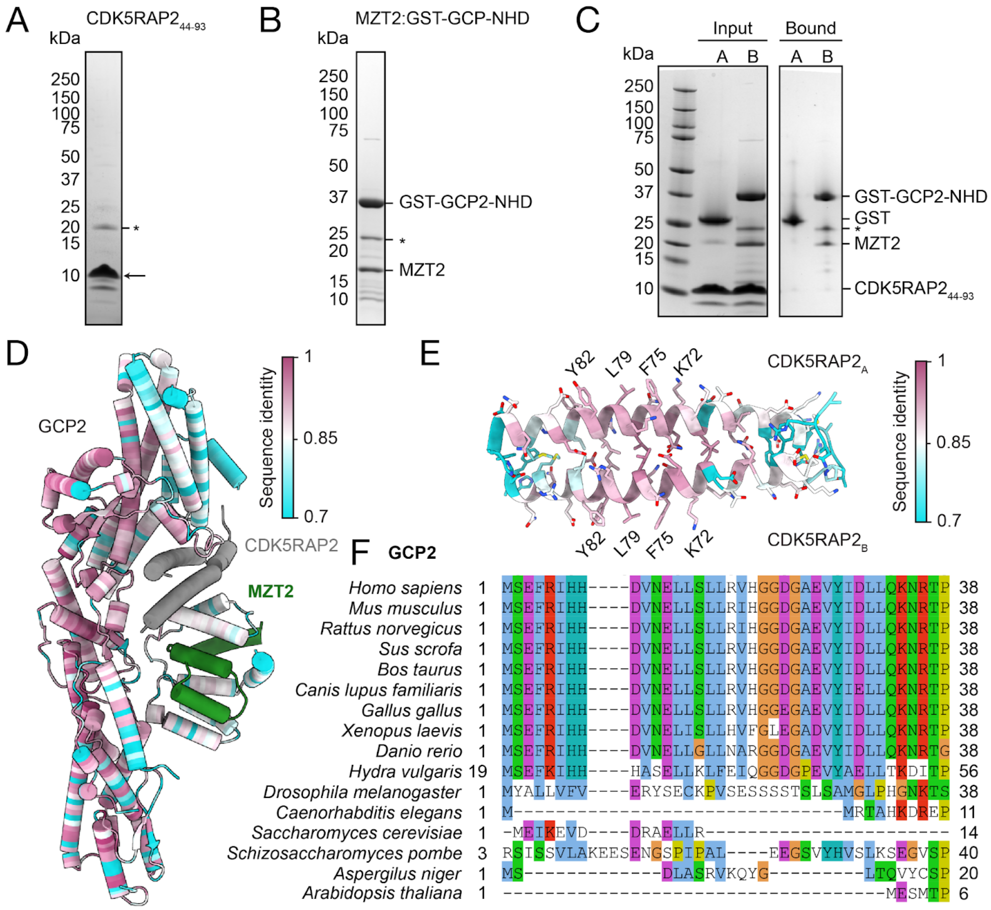


**Figure S7. CDK5RAP2$_{44\text{-}93}$ and MZT2:GCP2-NHD do not interact in the absence of additional γ-TuRC interfaces.** A)-B) Coomassie-stained SDS-PAGE gels (4-20% Tris-glycine) of CDK5RAP2$_{44\text{-}93}$ (A) and MZT2:GST-GCP-NHD. Arrow in A) points to the CDK5RAP2$_{44\text{-}93}$ construct, while asterisks in A)-B) indicate a contaminant. The predicted molecular weight of CDK5RAP2$_{44\text{-}93}$ after cleavage is 7.3 kDa; that the band runs higher than 10 kDa may be due to difficulty in disrupting dimerization under SDS-PAGE conditions. C) Coomassie-stained SDS-PAGE (10% Tris-tricine) of input reactions (left) and proteins bound to glutathione beads after washing (right). The CDK5RAP2$_{44\text{-}93}$ concentration was 2 μM (dimer), and the GST (A lanes) and GST-GCP2-NHD (B lanes) concentrations were 5 μM. Under these conditions CDK5RAP2$_{44\text{-}93}$ is not detected in the bound fractions. Asterisk in C) indicates the same contaminating protein band in the MZT2:GST-GCP2-NHD (panel B). D) Relative amino acid sequence conservation of human GCP2 judged by the multiple sequence alignment of 715 non-redundant sequences retrieved from UniProt plotted on a GCP2 at position 7. GCP2 residues are colored according to sequence identity in MSA, most conserved positions are shown in maroon, least conserved are colored cyan, MZT2 is shown in green and two copies of CDK5RAP2$_{44\text{-}93}$ are grey. E) Relative amino acid sequence conservation of CDK5RAP2$_{44\text{-}93}$ assessed from a multiple sequence alignment of 709 non-redundant sequences from UniProt. Color code covers the same range of sequence identity as in (D). Residues of the highest conservation and greatest contribution to the binding pocket are highlighted. F) Multiple sequence alignment of GCP2 proteins from various model organisms. Residues corresponding to human GCP2 1-38 fragments are shown. Residues are colored using the default Clustal color code in JalView [6,7].

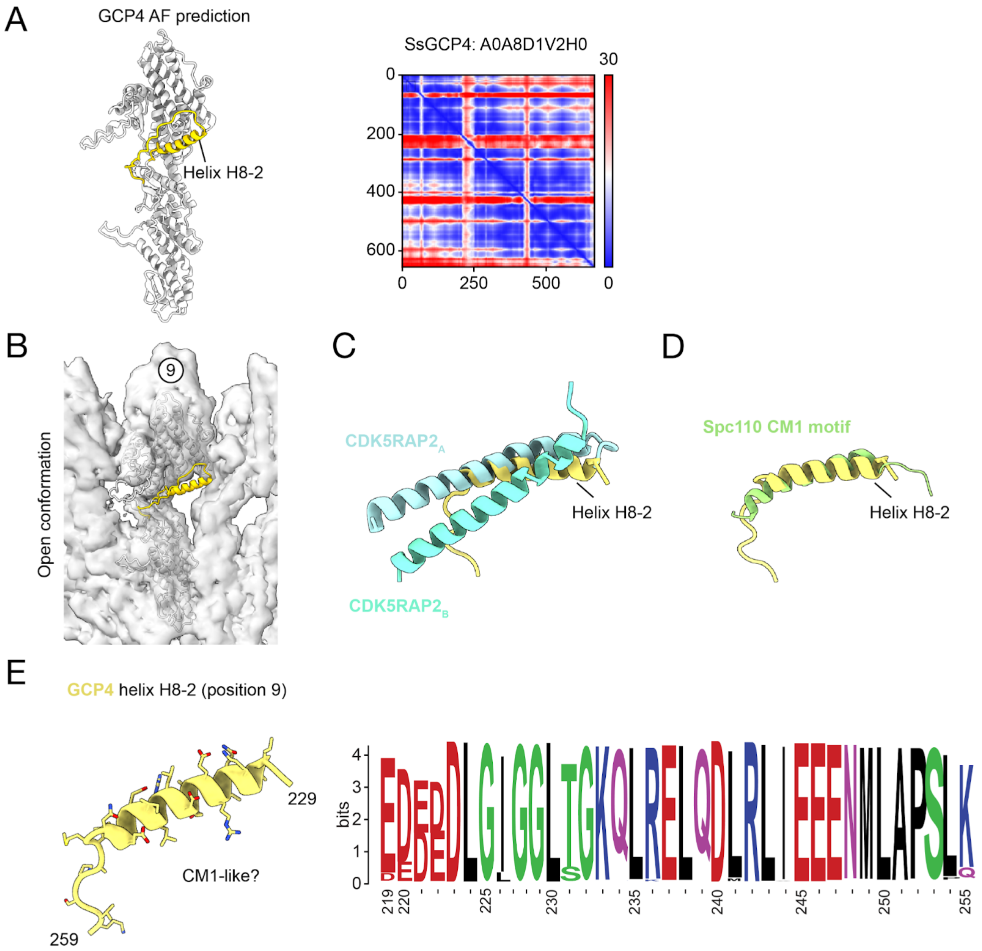


**Figure S8. Supplementary information for GCP4 helix H8-2.** A) Cartoon representation of an AlphaFold prediction for full-length *S. scrofa* GCP4. Helix H8-2 is coloured in yellow. The PAE plot is shown on the right. B) The model in A) rigid body-fitted into the open γ-TuRC conformation density map at position 9 (transparent surface). C) Cartoon representation of the CDK5RAP2 dimer and helix H8-2 from. Overlay generated by aligning GCP2 from position 7 of the CMG-decorated conformation onto GCP4 at position 9. D) Same principle as in C), but aligning Spc97 and displaying the Spc110 CM1 motif on top of helix H8-2. E) Left: Cartoon and stick representation of GCP4 helix H8-2. Right: Conservation of the helix H8-2 sequence in GCP4s across species. Generated using WebLogo [8].

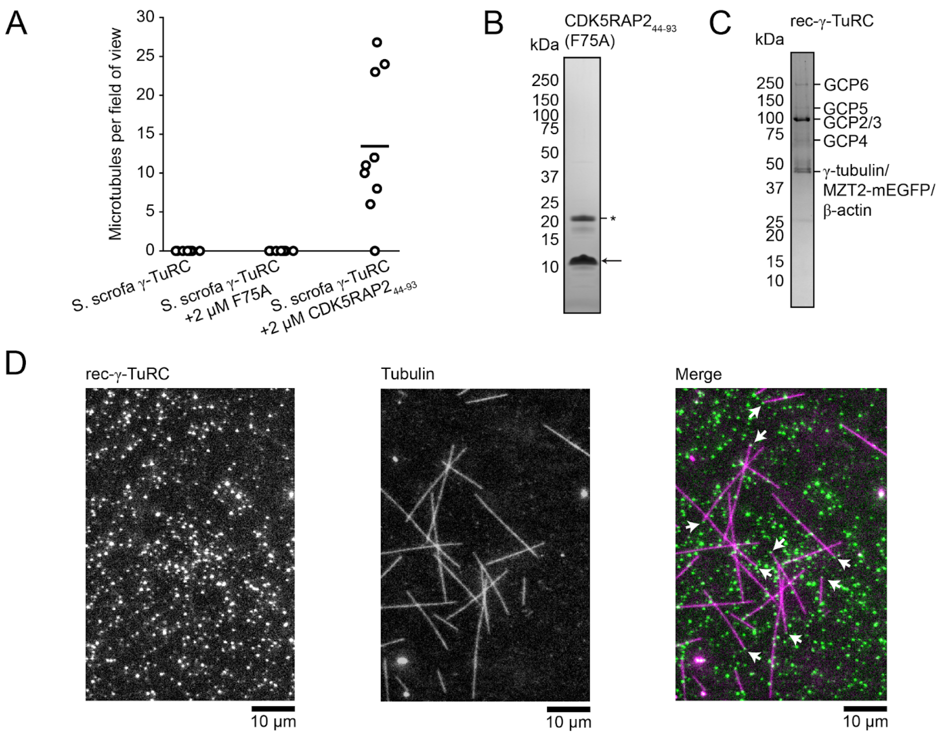


**Figure S9. Supplementary information for TIRF assays.** A) Plot of the number of microtubules detected per field of view for non-specifically adsorbed *S. scrofa* γ-TuRC without added $CDK5RAP_{44\text{-}93}$ construct (left), in the presence of 2 uM of the F75A $CDK5RAP_{44\text{-}93}$ mutant (middle) and 2 uM of WT $CDK5RAP_{44\text{-}93}$ (right; n = 9 fields of view from 3 independent experiments per condition). B)-C) Coomassie-stained SDS-PAGE gels (4-20% Tris-glycine) of $CDK5RAP2_{44\text{-}93}$ with the F75A mutation (B) and rec-γ-TuRC (C). Arrows and asterisks in B) point to the $CDK5RAP2_{44\text{-}93}$ construct and a contaminant, respectively. As for WT, the predicted molecular weight of $CDK5RAP2_{44\text{-}93}$ F75A after cleavage is 7.3 kDa; that the band runs higher than 10 kDa may be due to difficulty in disrupting dimerization under SDS-PAGE conditions. For both WT and F75A, LC-MS/MS was used to confirm the presence of CDK5RAP2 as the dominant source of tryptic peptides (data not shown). D) TIRF microscopy assay micrographs showing the rec-γ-TuRC channel (left), the tubulin channel (middle), and the merge (right). Microtubule ends co-localizing with rec-γ-TuRC are indicated by white arrows.

### Supplementary Videos

**Video S1. Overview of the open to closed transition in the CMG-decorated γ-TuRC.** The beginning of the movie depicts a cartoon representation of the S. scrofa γ-TuRC in the open conformation morphing into the CMG-decorated conformation, emphasizing the locations of CMG decorations and both GCP4 helices H8-2. Afterwards, these decorations remain highlighted in cartoon representation against the rest of the γ-TuRC, which is shown as a surface representation. At the end of the movie, the surface representation is removed to allow the video to loop.

**Supplementary Tables**

**Table S1.** Mass spectrometry analysis of *S. scrofa* brain γ-TuRC.

| Component | Accession reported by LC-MS/MS | Peptides (unique/ total) | Coverage | iBAQ ratio relative to GCP6 | Accession used for model building | Model molecular weight in kDa** |
|---|---|---|---|---|---|---|
| γ-tubulin | F2Z562 | 6/27 | 68% | 33.8 | A0A287BRH5 | 51.1 |
| GCP2 | F1SCV1 | 42/42 | 51% | 12.8 | A0A480VJI0 | 102.5 |
| GCP3 | F1RN46 | 47/47 | 47% | 11.6 | F1RN46 | 103.0 |
| GCP4 | F1SI61 | 23/23 | 49% | 2.8 | A0A8D1V2H0 | 76.0 |
| GCP5 | I3L738 | 16/16 | 18% | 0.7 | A0A287B1K1 | 119.9 |
| GCP6 | F1RXS1 | 22/22 | 16% | 1 | A0A8W4FDV6 | 188.4 |
| MZT1 | A0A287AS02 | 2/2 | 37% | 2.3 | A0A287AS02 | 8.3 |
| MZT2 | A0A287AYE6 | 6/6 | 37% | 7.2 | F1RK97 | 15.9 |
| β-actin | Q6QAQ1 | 1/19 | 57% | 10.8 | Q6QAQ1 | 41.7 |
| CDK5RAP2* | I3LKY1 | 7/7 | 4% | 39.8 | CDK5RAP2* | 4.5 |
| β-spectrin | A0A287BDN2 | 39/41 | 21% | 0.5 | n.a. | 274.6 |

* = Human sequence used for $CDK5RAP2_{51\text{-}100}$
** = Refers to sequences used for model building, except for β-spectrin
n.a. = not applicable

**Table S2.** Cryo-EM data collection.

| | **Dataset 1** | **Dataset 2** | **Dataset 3** | **Dataset 4** | **Dataset 5** | **Dataset 6** |
|---|---|---|---|---|---|---|
| Microscope | Titan Krios | Titan Krios | Titan Krios | Titan Krios | Titan Krios | Titan Krios |
| Voltage (kV) | 300 | 300 | 300 | 300 | 300 | 300 |
| Magnification | 81,000 X | 81,000 X | 81,000 X | 81,000 X | 81,000 X | 130,000 X |
| Pixel size (Å/px) | 1.06 | 1.06 | 1.06 | 1.06 | 1.06 | 0.64 |
| Camera | K3 | K3 | K3 | K3 | K3 | K3 |
| GIF slit | 20 eV | 20 eV | 20 eV | 20 eV | 15 eV | 20 eV |
| Mode | CDS | CDS | CDS | CDS | CDS | CDS |
| Software | EPU | EPU | EPU | EPU | EPU | EPU |
| Spot size | 5 | 5 | 6 | 6 | 7 | 6 |
| Beam diameter | 1.25 | 1.29 | 0.89 | 1.07 | 0.98 | 0.88 |
| # of grids | 1 | 2 | 1 | 2 | 1 | 1 |
| # of movies | 5,256 | 29,548 | 15,561 | 9,595 | 22,218 | 27,546 |
| # of frames | 40 | 40 | 40 | 40 | 60 | 40 |
| Exposure time (s) | 2 | 2 | 1 | 2 | 1 | 1 |
| Dose Rate ($e^-$/px/s) | 7.68 | 7.5 | 7.5 | 7.58 | 7.87 | 7.17 |
| Total exposure ($e^-/Å^2$) | 50 | 52 | 51 | 51 | 78 | 61 |
| Defocus range (μm) | -1.6/-3 | -1.6/-3 | -1.4/-2.8 | -0.8/-2.8 | -0.8/-2.5 | -0.8/-2.4 |

**Table S3.** Cryo-EM data processing statistics.

| | Consensus map | Overall (open) | Pos. 1 to 3 (open) | Pos. 3 to 11 (open) | Pos. 10 to 12 (open) | Partially-assembled | Overall (CMG-dec.) | Pos. 1 to 3 (CMG-dec.) | Pos. 1 to 8 (CMG-dec.) | Pos. 9 to 11 (CMG-dec.) | Pos. 13 (CMG-dec.) | Pos. 14 (CMG-dec.) |
|---|---|---|---|---|---|---|---|---|---|---|---|---|
| Processing pixel size (Å/px) | 1.4133 | 1.4133 | 1.4133 | 1.4133 | 1.4133 | 1.4133 | 1.4133 | 1.4133 | 1.4133 | 1.4133 | 1.4133 | 1.4133 |
| Box size (pix) | 384 | 384 | 384 | 384 | 384 | 384 | 384 | 384 | 384 | 384 | 384 | 384 |
| Symmetry imposed | C1 | C1 | C1 | C1 | C1 | C1 | C1 | C1 | C1 | C1 | C1 | C1 |
| Number of particles | 456,707 | 114,154 | 148,619 | 183,770 | 49,860 | 22,302 | 21,540 | 21,540 | 25,580 | 74,139 | 28,818 | 65,806 |
| Resolution (Å) (FSC 0.143 cutoff) | 4.1 | 4.6 | 4.7 | 4.1 | 5.2 | 6.2 | 7.5 | 7.5 | 6.5 | 6.5 | 7.5 | 6.2 |
| Resolution range (Å) (ResMap) | (3 - 10) | (3 - 10) | NC | NC | NC | (3 - 10) | (3.5 - 12) | NC | NC | NC | NC | NC |
| Model building | No | Yes | Yes | Yes | Yes | Yes | Yes | Yes | Yes | Yes | Yes | Yes |

*NC, Not calculated

**Table S4.** Model building and refinement statistics

| | Overall (open) | Pos. 1 to 3 (open) | Pos. 3 to 11 (open) | Pos. 10 to 12 (open) | Partially-assembled | Overall (CMG-dec.) | Pos. 1 to 3 (CMG-dec.) | Pos. 1 to 8 (CMG-dec.) | Pos. 9 to 11 (CMG-dec.) | Pos. 13 (CMG-dec.) | Pos. 14 (CMG-dec.) |
|---|---|---|---|---|---|---|---|---|---|---|---|
| Initial models | AF, HM | AF, HM | AF, HM | AF, HM | AF, HM | AF, HM | AF, HM | AF, HM | AF, HM | AF, HM | AF, HM |
| Model-to-Map fit | 0.6537 | 0.6962 | 0.7672 | 0.5707 | 0.8048 | 0.7360 | 0.7220 | 0.7619 | 0.6100 | 0.7034 | 0.2672 |
| Number of models | 1 | 1 | 1 | 1 | 1 | 1 | 1 | 1 | 1 | 1 | 1 |
| Number of Chains | 34 | 6 | 21 | 6 | 10 | 46 | 15 | 32 | 6 | 11 | 11 |
| Number of Residues | 15,602 | 3,210 | 9,901 | 3,286 | 4,440 | 16,662 | 4,780 | 10,021 | 3,137 | 4,574 | 3,652 |
| All-atom Clashscore | 20.20 | 17.76 | 13.59 | 18.92 | 16.49 | 16.14 | 18.65 | 21.94 | 18.39 | 17.03 | 16.88 |
| Outliers (%) | 0.12 | 0.09 | 0.03 | 0.15 | 0.09 | 0.08 | 0.04 | 0.07 | 0.10 | 0.35 | 0.16 |
| Allowed (%) | 4.26 | 3.46 | 3.15 | 5.63 | 3.94 | 3.72 | 3.77 | 4.05 | 4.34 | 4.61 | 4.98 |
| Favored (%) | 95.62 | 96.45 | 96.82 | 94.22 | 95.97 | 96.20 | 96.19 | 95.88 | 95.57 | 95.04 | 94.85 |
| Rotamer outliers (%) | 0.01 | 0.00 | 0.00 | 0.07 | 0.00 | 0.01 | 0.00 | 0.01 | 0.00 | 0.00 | 0.00 |
| $C_\beta$ deviations (%) | 0.00 | 0.00 | 0.00 | 0.00 | 0.00 | 0.00 | 0.00 | 0.01 | 0.00 | 0.00 | 0 |
| Bond length (Å) | 0.004 | 0.003 | 0.004 | 0.004 | 0.003 | 0.006 | 0.011 | 0.008 | 0.003 | 0.003 | 0.010 |
| Bond angles (º) | 0.660 | 0.637 | 0.614 | 0.769 | 0.604 | 0.602 | 0.640 | 0.676 | 0.585 | 0.654 | 0.686 |
| EMDB | EMD-XXXX | EMD-XXXX | EMD-XXXX | EMD-XXXX | EMD-XXXX | EMD-XXXX | EMD-XXXX | EMD-XXXX | EMD-XXXX | EMD-XXXX | EMD-XXXX |
| PDB | PDB-XXXX | | | | PDB-XXXX | PDB-XXXX | | | | | |
